# Global analysis of putative phospholipases in the malaria parasite *Plasmodium falciparum* reveals critical factors for parasite proliferation

**DOI:** 10.1101/2021.06.28.450158

**Authors:** Paul-Christian Burda, Abhinay Ramaprasad, Emma Pietsch, Sabrina Bielfeld, Christoph Söhnchen, Louisa Wilcke, Jan Strauss, Dominik Schwudke, Aaron Sait, Lucy M Collinson, Michael J Blackman, Tim-Wolf Gilberger

## Abstract

For its replication within red blood cells, the malaria parasite is highly dependent on correctly regulated lipid metabolism. Enzymes involved in lipid metabolic processes are therefore potential drug targets. We here provide a functional analysis of the 20 putative phospholipases that are expressed by asexual blood stages of *Plasmodium falciparum*. We reveal a high level of redundancy among members of this group, but using conditional mislocalization and gene disruption techniques we show that the phosphoinositide-specific phospholipase C (PF3D7_1013500) has a previously unrecognized essential role in intracellular parasite maturation. In addition, we demonstrate that the patatin-like phospholipase PF3D7_1358000 localizes to the mitochondrion. Parasites lacking this enzyme display a severe growth phenotype and defects in mitochondrial morphogenesis and function leading to hypersensitivity towards proguanil and inhibitors of the mitochondrial electron transport chain including atovaquone. This demonstrates that regulated mitochondrial lipid homeostasis is necessary for mitochondrial function and coordinated division during parasite multiplication.

## INTRODUCTION

With an estimated 228 million cases per year worldwide and more than 400,000 deaths, malaria remains one of the most important human health threats (WHO, 2019). The replication of protozoan parasites of the genus *Plasmodium* within red blood cells (RBCs) and the associated transformation and destruction of these cells are responsible for the clinical symptoms of the disease. With no licensed vaccine widely available and widespread resistance of the parasite to available drugs, there is an urgent need to better understand the biology of the malaria parasite in order to identify suitable targets for new intervention strategies.

Malaria parasites are transmitted by the bite of an infected *Anophele*s mosquito and initially establish in their vertebrate host by multiplying in hepatocytes. From here, parasites are released into the bloodstream, where they undergo repeated cycles of replication within RBCs (reviewed in (De Niz et al., 2017)). Central to intraerythrocytic growth of parasites is an intense period of membrane biogenesis. Not only do the intracellular parasites need to extend the parasite plasma membrane and replicate their organelles during the formation of daughter cells, they also have to support expansion and maturation of the parasitophorous vacuole membrane (PVM), which surrounds them during their multiplication. As a consequence of this, the phospholipid content of the RBC increases almost fivefold during intraerythrocytic development (Tran et al., 2016). Fatty acids, the building blocks of lipids, are largely taken up from the host, but due to the presence of a functional FASII system in the apicoplast, a non-photosynthetic plastid derived from algae, the parasite can also synthesize fatty acids *de novo*; this is particularly important for parasite development in the liver (reviewed in (Tarun et al., 2009)). Generation of membranes not only requires *de novo* synthesis and acquisition but also the degradation of phospholipids, a function performed by phospholipases. These enzymes hydrolyze specific ester bonds in phospholipids and are classified into four groups, A, B, C, and D based on their hydrolysis activity (reviewed in (Flammersfeld et al., 2017)). Although phospholipases likely play key functions for *Plasmodium* cell biology, little is known about their role in proliferation of the malaria parasite and information regarding phospholipase essentiality is incomplete.

We here have performed a comprehensive functional analysis of the phospholipase gene family in the most virulent malaria parasite species *Plasmodium falciparum* during its asexual multiplication within RBCs. Using conditional inactivation techniques, we provide evidence for a physiological function of the phosphoinositide-specific phospholipase C (PI-PLC) during intracellular parasite maturation, long before its previously perceived role at parasite egress and invasion. We additionally show that, during their development within RBCs, parasites express a patatin-like phospholipase that regulates mitochondrial morphogenesis and function, representing a novel role for this class of enzymes in protozoan parasites.

## RESULTS

### Gene deletion screening of the *Plasmodium* phospholipase family in asexual blood stages

We started to systematically investigate the function of *Plasmodium* phospholipases in *P. falciparum* asexual blood stages by first searching the *Plasmodium* genome for genes encoding proteins containing putative lipase/phospholipase-related domains (Plasmodb.org, (Aurrecoechea et al., 2009)). This resulted in a list of 27 genes encoding enzymes with putative phospholipase function (Figure 1 – figure supplement 1) that also included the putative phospholipases identified previously (Flammersfeld et al., 2017). For 20 of these 27 genes there exists mass spectrometric evidence for expression in asexual blood stage parasites (Bowyer et al., 2011; Cobbold et al., 2016; Florens et al., 2002, 2004; Lasonder et al., 2012, 2015; Oehring et al., 2012; Pease et al., 2013; Silvestrini et al., 2010; Solyakov et al., 2011; Treeck et al., 2011). We therefore focused subsequent efforts on these 20 genes and characterized their essentiality for the erythrocytic parasite life cycle.

For this, we performed targeted gene disruption (TGD) using the selection-linked integration (SLI)-system (Birnbaum et al., 2017) (Figure 1A). Of the 20 transfected targeting constructs, each designed to disrupt expression of the targeted gene, we obtained outgrowth of viable parasites displaying correct integration into their respective gene loci in 15 cases, indicating that the corresponding genes are not essential for *in vitro* parasite growth (Figure 1B, Figure 1 – figure supplement 2). For the remaining five putative phospholipase genes we consistently failed to obtain viable parasites harboring correctly integrated targeting plasmid, suggesting that these genes are important or essential for propagation of asexual blood stage parasites (Figure 1B). Analysis of the obtained 15 mutant lines for potential growth defects revealed that only the patatin-like phospholipase PF3D7_1358000 mutant consistently showed a reduction in growth rate of ∼50% in comparison to wild type (WT) parasites over two parasite cycles (Figure 1C, Figure 1 – figure supplement 3). All the other mutant parasite lines displayed no or very slightly reduced growth rates. These data are overall in agreement with a recent genome-wide saturation mutagenesis screen in *P. falciparum* (Zhang et al., 2018). However, there were several exceptions to this. On the one hand, the patatin-like phospholipase 1 (PF3D7_0209100), for which we could not obtain transgenic knockout (KO) parasites, was identified as being nonessential by Zhang *et al*. and the redundant function for asexual blood stage development was recently confirmed by two independent groups (Flammersfeld et al., 2019; Singh et al., 2019). On the other hand, five of the analyzed genes suggested to be essential by Zhang *et al*. could all be inactivated by our SLI-system, underlining the need to verify global-scale screening data at the single gene level. Another large-scale knockout screen was performed in blood stages of the rodent malaria model *Plasmodium berghei* (Bushell et al., 2017). Interestingly, of the eight putative phospholipase orthologs analyzed in that screen only the genes encoding PI-PLC and the patatin-like phospholipase PF3D7_1358000 could not be disrupted (Figure 1B), consistent with our new data and supporting an important role for these two enzymes in intraerythrocytic parasite replication.

**Figure 1.**
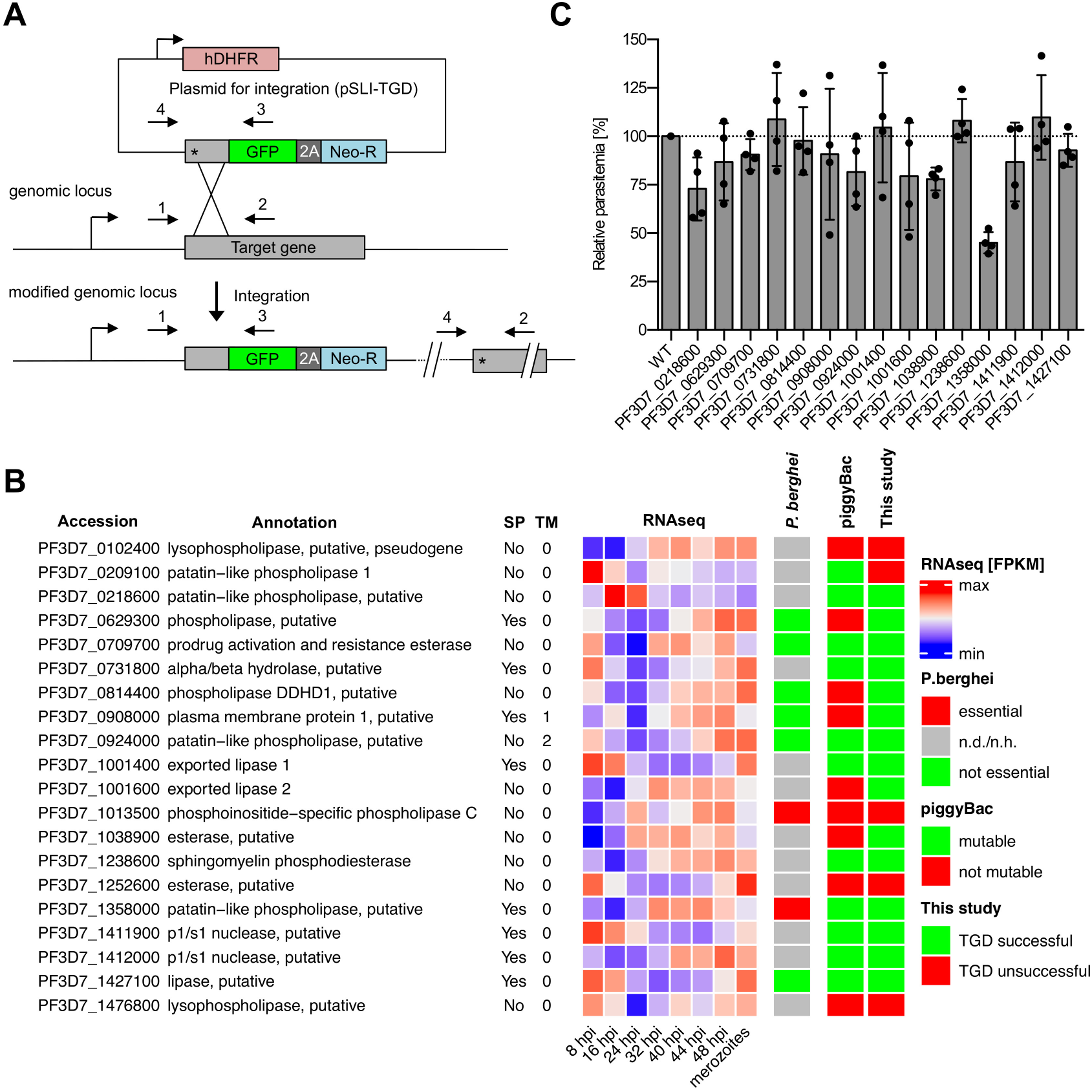
Gene disruption screen of the 20 predicted *P. falciparum* phospholipases expressed during blood stage development. A) Schematic of the selection-linked-integration (SLI) strategy used for targeted gene disruption (TGD)-based essentiality screening of the 20 putative phospholipases that show expression evidence in blood stages by mass spectrometry. Localization of primers used to detect successful integration of targeting constructs by PCR are indicated. Integration PCR results are displayed in Figure 1-figure supplement 2. 2A, skip peptide; Neo-R, neomycin-resistance gene; asterisks, stop codons; arrows, promoters. B) Results of the gene deletion screen compared to the results of genome-wide knockout screens in *P. falciparum* using piggyBack transposon-based mutagenesis (Zhang et al., 2018) and in *P. berghei* (Bushell et al., 2017), respectively. RNAseq expression data is derived from (Wichers et al., 2019). SP, signal peptide; TM, transmembrane domain. For further details see Figure 1 – figure supplement 1. C) Flow cytometry-based growth analysis of synchronous phospholipase mutant parasite lines after two erythrocytic cycles in comparison to 3D7 WT parasites. Relative growth of each parasite line is shown in comparison to 3D7 WT parasites, the growth rate of which was normalized to 100% in each experiment. Shown are means +/- SD of four independent growth experiments per parasite line. Raw parasitemia values are shown in Figure 1 – figure supplement 3.

### PI-PLC is essential for parasite proliferation

Based on our SLI-based gene disruption data indicating that the single *P. falciparum* PI-PLC (PF3D7_1013500) is critical for parasite growth (Figure 1B), we decided to further investigate the functional role of this putative enzyme. *P. falciparum* PI-PLC is 1,385 amino acids in length and contains all the functional domains typical for PI-PLC enzymes of the delta subclass, including: i) a lipid binding pleckstrin homology (PH)- domain (residues 80-209); ii) a calcium-binding EF-hand motif (residues 217-304); iii) a catalytic domain consisting of an X- (residues 624-769) and Y-domain (residues 972-1087); and iv) a calcium/lipid-binding C2 domain (residues 1279-1383) (Figure 2A) (Raabe et al., 2011a). To analyze the subcellular localization of PI-PLC and study its function, we made use of the recently developed conditional knocksideways system (Birnbaum et al., 2017). For this, we first tagged the endogenous PI-PLC coding sequence by generating a C-terminal fusion to GFP flanked by two FKBP-domains (Figure 2 – figure supplement 1). We then expressed in this parasite line a ‘mislocalizer’ protein called NLS-ML, consisting of mCherry fused to an FRB-domain and a nuclear localization signal. Addition of the small molecule Rapalog (Rapa) mediates heterodimerization of the NLS-ML and PI-PLC-GFP-FKBP proteins, removing the latter from its physiological site of action to the nucleus (Birnbaum et al., 2017). The resulting parasite line, called PI-PLC-GFP-knocksideways (PI-PLC-GFP-KS), was used for subsequent localization and functional characterization.

**Figure 2.**
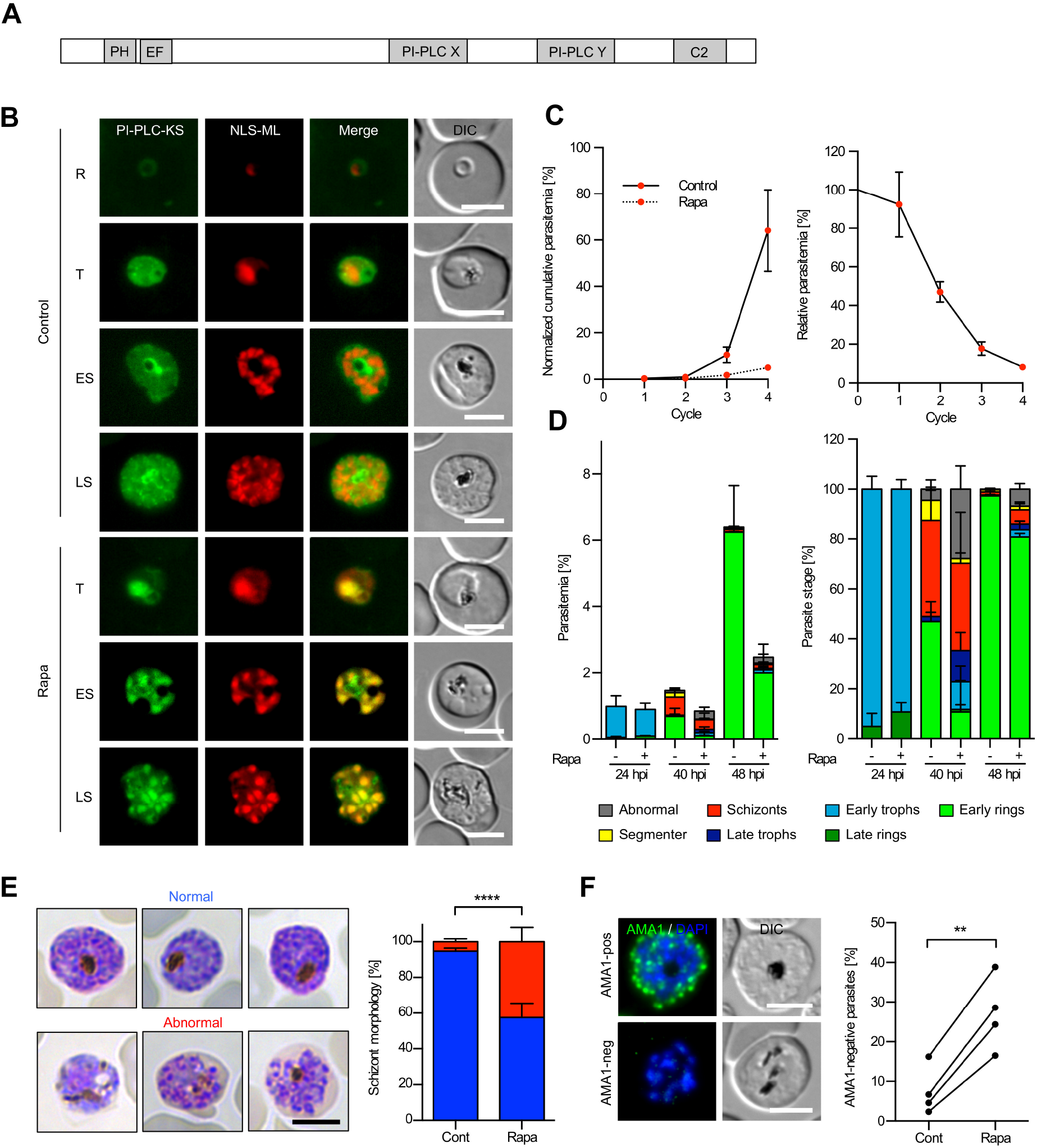
PI-PLC has an essential role in trophozoite and schizont development. A) Schematic representation of the functional domains of PI-PLC. B) Live cell microscopy of ring (R), trophozoite (T) early schizont (ES) and late schizont (LS) of PI-PLC-GFP-KS parasites expressing endogenously FKBP-GFP-tagged PI-PLC (green) in addition to a nuclear mislocalizer fused to mCherry (NLS-ML, red). Parasites were either untreated (control) or treated with Rapa to conditionally mislocalize the PI-PLC to the nucleus. DIC, differential interference contrast. C) Growth over four erythrocytic cycles of PI-PLC-GFP-KS parasites treated with Rapa in comparison to untreated control parasites as determined by flow cytometry. For calculation of the normalized cumulative parasitemia (shown on the left), parasitemia levels of untreated parasites were normalized to 1% in the second cycle. For calculation of relative parasitemia values (shown on the right), the parasitemia of Rapa-treated parasites was divided by the parasitemia of the respective untreated controls. Shown are means +/- SD of four independent experiments. D) Stage and parasitemia quantification of control and Rapa-treated PI-PLC-GFP-KS parasites at 24, 40 and 48 hpi. Shown are means +/- SD of four independent experiments. E, F) Schizont morphology and AMA1 expression of untreated and Rapa-treated PI-PLC-GFP-KS schizonts at 48 hpi, which were cultured in presence of C2 for 8 hours to prevent egress. In (E) schizont morphology was determined by microscopic examination of Giemsa-stained parasites. Shown are means +/- SD of four independent experiments, in which a total of 156 control and 153 Rapa-treated schizonts were analyzed. Statistical evaluation was by unpaired students t-test (**** p < 0.0001). Representative images of normal and abnormal schizonts are shown on the left. In (F) AMA1-expression was assessed by IFA. Shown are means +/- SD of four independent experiments, in which a total of 306 control and 356 Rapa-treated schizonts were analyzed. Statistical analysis was by a paired students t-test (** p < 0.01). Representative AMA1-positive and AMA1-negative schizonts are shown on the left. All scale bars, 5 µm.

Previous RNAseq studies have shown peak expression of the *pi-plc* gene during trophozoite development (López-Barragán et al., 2011). In accord with this, live microscopic examination of untreated PI-PLC-GFP-KS parasites revealed a GFP signal in trophozoite and schizont stage parasites that was mainly confined to the parasite cytoplasm. Interestingly, in mature schizonts the signal appeared to partially surround developing daughter merozoites, suggesting a potential association of PI-PLC with the parasite plasma membrane (Figure 2B, upper panels). Treatment of synchronous ring stage PI-PLC-GFP-KS parasites with Rapa led to rapid redistribution of the PI-PLC signal to the nucleus, as expected, leading to efficient colocalization with the NLS-ML signal (Figure 2B, lower panels). To investigate the effects of this conditional PI-PLC mislocalization on parasite development, we compared the replication rates of untreated and Rapa-treated parasites over four erythrocytic cycles using flow cytometry. This showed that the Rapa-treated parasites displayed an ∼50% reduced multiplication rate per cycle, leading to a reduction in replication of more than 90% after four erythrocytic cycles (Figure 2C). The residual multiplication capacity of the Rapa-treated PI-PLC-GFP-KS parasites is likely explained by the fact that only about 90 +/- 5% (mean +/- SD of three independent quantification experiments, total n=356) of these parasites expressed the NLS-ML construct. This substantial reduction in parasite replication upon conditional mislocalization of PI-PLC is consistent with the results of our SLI-based gene disruption screen (Figure 1), confirming that PI-PLC indeed plays an important role in erythrocytic parasite growth.

### PI-PLC is involved in trophozoite and schizont development

To determine the specific stage(s) in the erythrocytic developmental cycle affected by conditional mislocalization of PI-PLC, we monitored the development of tightly synchronized control and Rapa-treated PI-PLC-GFP-KS parasites by light microscopic examination of Giemsa-stained thin blood films. While parasite development appeared to be unaffected over the first 24 hours post RBC invasion (24 hpi), clear effects on parasite maturation were detectable in Rapa-treated PI-PLC-GFP-KS parasites by 40 and 48 hpi (Figure 2D). At 40 hpi, ∼25% of Rapa-treated parasites were still at the trophozoite stage, in contrast to the untreated parasites at this time point, in which hardly any trophozoites were detectable. Furthermore and in contrast to control parasites, ∼40% of the Rapa-treated parasites that formed schizonts displayed abnormal morphology. Together, these observations suggest that PI-PLC is involved in trophozoite and schizont development. Likely as a consequence of this, ring stage parasitemia values at 40 and 48 hpi were reduced by more than 60% in the Rapa-treated parasites (Figure 2D).

To analyze in further detail this potential function of PI-PLC during schizont development, we used an inhibitor of the parasite cGMP-dependent protein kinase G (PKG), called compound 2 (C2, which prevents egress) to synchronize parasites at mature schizont stage (Taylor et al., 2010). Examination of these C2-arrested schizonts revealed that more than 40% of Rapa-treated PI-PLC-GFP-KS parasites were dysmorphic (Figure 2E). In line with this, analysis by immunofluorescence assay (IFA) of the C2-arrested parasites showed that a high proportion of Rapa-treated parasites failed to express the late stage specific marker AMA1 (Figure 2F). Collectively, these results highlight a crucial role for PI-PLC in intraerythrocytic parasite maturation.

To test whether the maturation phenotype upon conditional inactivation of PI-PLC is associated with a perturbation of the lipid homeostasis, we subjected untreated and Rapa-treated PI-PLCK-GFP-KS trophozoites (30 hpi) and schizonts (40 hpi) to lipidomic analysis. In this semi-targeted lipid analysis, 266 lipids were quantified covering 15 lipid classes and cholesterol. In line with the maturation phenotype, we observed that Rapa-treated trophozoites and schizonts had a significantly reduced lipid content in comparison to untreated parasites. Apart from this, only minor alterations in the lipid profile were observed, including a reduced absolute concentration of phosphatidylglycerol and cardiolipin per schizont (Figure 2 – figure supplement 2, Supplementary file 2). Levels of diacylglycerol, a primary metabolite of PI-PLC activity, were unchanged, suggesting an only minor contribution of PI-PLC activity to the total intracellular pool of diacylglycerols.

### Conditional disruption of PI-PLC confirms its essentiality for *P. falciparum* asexual blood stage growth

The knocksideways system is a powerful tool to study the function of essential proteins that do not enter the secretory pathway (Birnbaum et al., 2017). However, under conditions where mislocalization is not 100% efficient, varying amounts of target protein can remain at the site of action and therefore functional. We therefore decided to further probe the function of PI-PLC using the rapamycin (RAP)-inducible dimerizable Cre recombinase (DiCre) system (Collins et al., 2013; Jones et al., 2016) to perform conditional disruption of the *pi-plc* gene. For this, a 3’-proximal segment of the *pi-plc* open reading frame encoding the predicted catalytic core of PI-PLC (the predicted X and Y domains) as well as the calcium/lipid-binding C2 domain, was targeted for excision by replacing the endogenous gene segment with a synthetic modified version using Cas9-enhanced homologous recombination. The modified sequence incorporated: i) a short synthetic intron containing a *loxP* site (loxPint) upstream of the catalytic domains; ii) the recodonized version of the segment encoding the WT amino acid sequence but with altered codon usage; iii) a C-terminal triple-hemagglutinin (3HA) epitope tag just preceding the translational stop codon; and iv) a second *loxP* site immediately following the translational stop codon (Figure 3A). The genetic modification was performed in the B11 *P. falciparum* line (Perrin et al., 2018), which stably expresses DiCre recombinase, a form of ‘split Cre’ which is activated by RAP-mediated heterodimerization. DiCre-mediated excision of the floxed sequence was expected to result in conditional inactivation of PI-PLC due to the deletion of its catalytic domains. The transgenic parasites (called PI-PLC:HA:loxPint) were cloned by limited dilution and two clonal parasite lines (D9 and F9) were isolated. The expected genetic modifications in both clones were confirmed by diagnostic PCR (Figure 3 – figure supplement 1). RAP treatment of tightly synchronized ring stage PI-PLC:HA:loxPint parasites resulted in the anticipated truncation of PI-PLC within the same erythrocytic cycle, as detected by PCR (Figure 3B), IFA (Figure 3C) and western blot (Figure 3D). To initially assess the viability of the resulting PI-PLC-null mutants, growth of RAP- and mock-treated cultures of the two PI-PLC:HA:loxPint clonal lines was monitored over the course of four erythrocytic cycles. PI-PLC-null parasites failed to proliferate, confirming our knocksideways-based indications that PI-PLC is crucial for viability during asexual blood stage replication of *P. falciparum* (Figure 3E). For more in-depth characterization, the F9 PI-PLC:HA:loxPint clone was used in all subsequent experiments.

**Figure 3.**
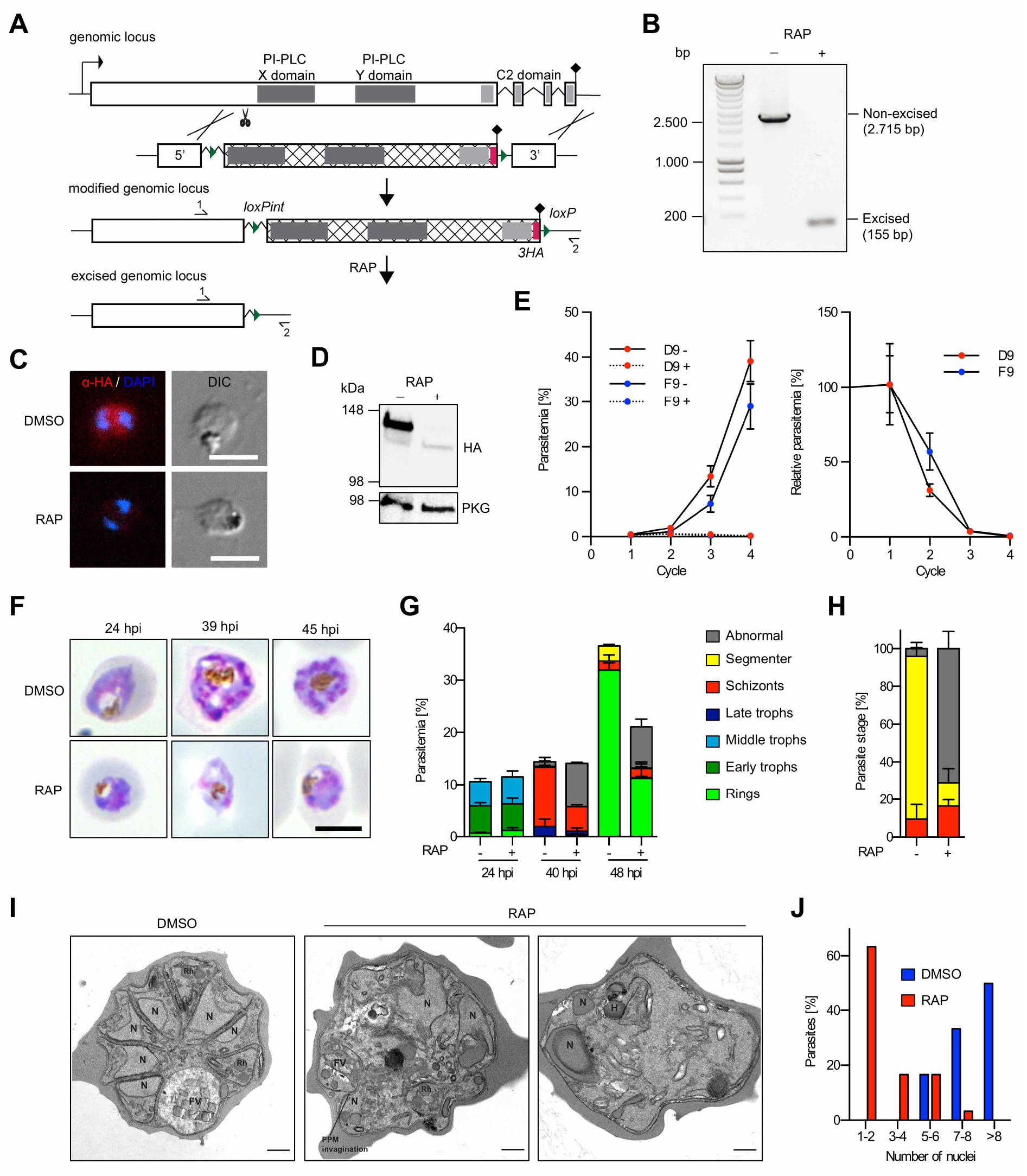
Conditional gene disruption confirms essentiality of PI-PLC. A) Schematic of the strategy used to generate a conditional PI-PLC knockout line (PI-PLC:HA:loxPint). The X and Y catalytic domains (dark grey) and the lipid-binding C2 domain (light grey) were floxed by introducing an upstream *loxP*-containing intron (loxPint) and a second *loxP* site downstream of the translational stop site (lollipop). Sites of targeted Cas9-mediated double-stranded DNA break (scissors), left and right homology arms for homology-directed repair (5’ and 3’), introduced *loxP* sites (arrow heads), recodonized sequences (hatched) and 3xHA epitope (red) are indicated. RAP-induced DiCre-mediated excision results in removal of the functional domains. Primers 1 and 2 (half arrows) were used for diagnostic PCR to assess excision. B) Confirmation of efficient gene excision by PCR. Samples were taken at 12 h post RAP- or mock (DMSO) treatment of ring stage parasites. Expected PCR amplicon sizes from non-excised and excised parasites are shown. Shown is one representative experiment (of five independent experiments). C) IFA images of mock or RAP-treated PI-PLC:HA:loxPint trophozoite stage parasites. Parasites were fixed at 33 hpi and stained with an anti-HA-antibody (red). DAPI-stained nuclei are shown in blue. DIC, differential interference contrast. Scale bars, 5 µm. D) Western blot of mature schizonts (45 hpi) showing successful RAP-induced ablation of PI-PLC-3xHA expression in the erythrocytic cycle of RAP addition. PKG served as a loading control. Note that PI-PLC-3xHA in DMSO-treated parasites runs slightly lower as compared to its calculated molecular weight of 167,4 kDa. E) Replication of mock- (solid line) and RAP-treated (dashed line) parasites from two clonal lines of PI-PLC:HA:loxPint over four erythrocytic cycles. For calculation of relative parasitemia values (shown on the right), the parasitemia of RAP-treated parasites was divided by the parasitemia of respective mock-treated control parasites. Shown are means +/- SD of three biological replicates (different blood sources). F) Giemsa-stained images of DMSO- and RAP-treated PI-PLC:HA:loxPint parasites at 24, 39 and 45 hpi. Representative images of two independent experiments are shown. Scale bars, 5 µm. G) Stage and parasitemia quantification of mock- and RAP-treated PI-PLC:HA:loxPint parasites at 24, 40 and 48 hpi. Shown are means +/- SD of three biological replicates. H, I, J) Morphological analysis of mock- or RAP-treated PLC:HA:loxPint parasites that were allowed to mature to egress-stalled schizonts from 46 to 49 hpi in the presence of C2. In (H) parasite morphology was assessed on Giemsa-stained blood smears. Shown are means +/- SD of three independent experiments. Color code same as in (G). In (I) and (J) parasite morphology was assessed using TEM. Representative images of mock- and RAP treated parasites are displayed in (I) and a quantification of nuclei is shown in (J). Results are representative of 18 DMSO- and 30 RAP-treated analyzed parasites. N, nucleus; FV, food vacuole; PPM, parasite plasma membrane; Rh, rhoptries; H, haemoglobin-containing cytostome. Scale bar, 500 nm.

Intracellular development of PI-PLC-null mutants within the erythrocytic cycle of RAP-treatment was studied by microscopic examination of Giemsa-stained parasites. This revealed that the PI-PLC-null mutants underwent apparently normal growth until late trophozoite stage, after which they developed morphological abnormalities with diffused nuclei and failed to progress through normal schizont maturation (Figure 3F,G). Schizont development was further analyzed by allowing mock- and RAP-treated parasites to reach maturity in the presence of the egress inhibitor C2. This revealed that >70% of PI-PLC-null parasites exhibited an abnormal morphology (Figure 3H). Analysis by transmission electron microscopy (TEM) of these C2-arrested parasites showed that more than 60% of PI-PLC-null parasites possessed poorly-defined subcellular organelles, and only 1-2 nuclei visible in the sections (rather than the 7 or more nuclei which were observed in about 80% of mock-treated control schizonts) (Figure 3I,J). Despite these developmental defects, haemoglobin-containing cytostomes and haemozoin crystals were evident in the digestive vacuole of the PI-PLC-null parasites, suggesting that the mutants retained the capacity to internalize and digest haemoglobin. Around 17% of PI-PLC-null parasites showed formation of 3-4 nuclei, well-formed rhoptries and parasite plasma membrane invaginations, pointing to the start of merozoite formation. However, we were unable to find more than a few well-segmented schizonts in the PI-PLC-null samples, in contrast to the majority of the mock-treated parasites, which formed well-segmented schizonts with clearly defined merozoites (Figure 3I,J). These microscopic observations show that most PI-PLC-null mutants show stunted growth in late trophozoite stages while a few of them develop to early schizont stage but stop short of becoming mature schizonts. Taken together, we concluded that lack of PI-PLC caused a severe growth defect during the trophozoite-schizont transition, suggesting that PI-PLC-mediated activity is critical for intraerythrocytic parasite development.

### A patatin-like phospholipase is critical for mitochondrial morphogenesis

Having established the importance of PI-PLC for parasite growth, we turned our attention to the putative phospholipase PF3D7_1358000, which was also indicated by our screen to be important for parasite replication. PF3D7_1358000 encodes a protein of 2,012 amino acids, containing a predicted signal peptide in addition to a patatin-like phospholipase (PNPLA) domain close to its C-terminal end (residues 1130-1405) (Figure 4A); based on this, the gene product is subsequently referred to as patatin-like phospholipase 2 (PNPLA2). Previous transcriptomic analyses indicate that peak expression of PNPLA2 in the asexual blood stage cycle occurs during schizont development (López-Barragán et al., 2011). In order to localize PNPLA2 in the parasite, we appended a C-terminal GFP-tag to the endogenous gene using the SLI system and confirmed the genetic modification by PCR (Figure 4 – figure supplement 1). Live fluorescence microscopy of PNPLA2-GFP parasites revealed that PNPLA2 localized to the mitochondrion, as shown by clear colocalization with MitoTracker Red (Figure 4B). In contrast, there was little or no colocalization with co-expressed mCherry directed to the apicoplast by fusion to the ACP-targeting sequence (Birnbaum et al., 2020), excluding localization of PNPLA2 to this organelle (Figure 4C).

**Figure 4.**
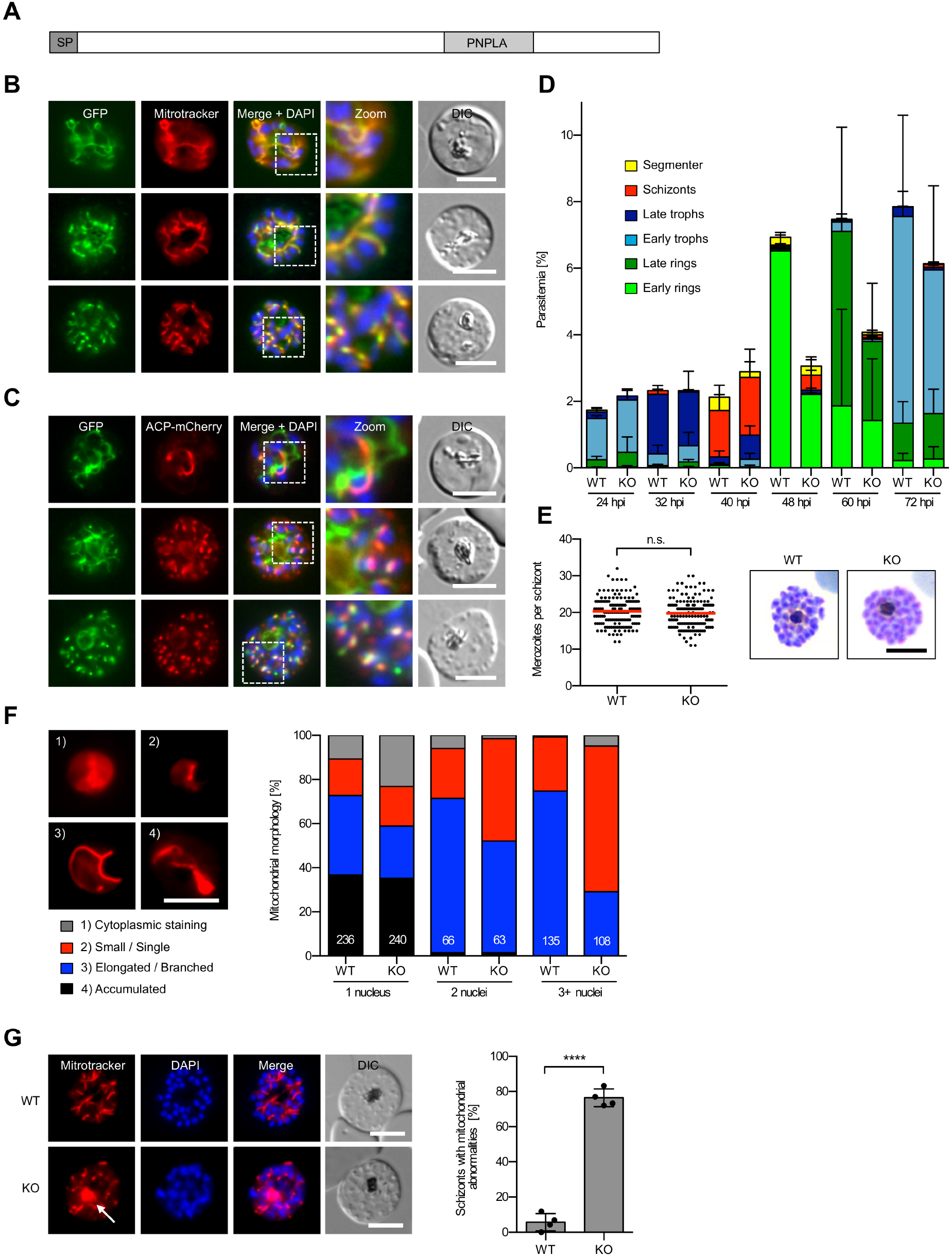
PNPLA2 localizes to the mitochondrion and is involved in mitochondrial morphogenesis. A) Schematic overview of the functional domains of PNPLA2. SP, signal peptide; PNPLA, patatin-like phospholipase domain. B, C) Live-cell microscopy of parasites expressing endogenously tagged PNPLA2-GFP (green). Parasites stained with MitoTracker Red (red) are shown in (B), whereas parasites co-expressing the apicoplast marker ACP-mCherry (red) are shown in (C). Merged images additionally contain DAPI-stained nuclei (blue). DIC, differential interference contrast. D) Stage quantification of WT and PNPLA2-KO parasites at different points after invasion. Shown are means +/- SD of three independent experiments. E) Merozoite numbers per schizont in WT and PNPLA2-KO parasites. Shown are pooled data of three independent experiments. In each experiment the number of merozoites per schizont was determined from 50 schizonts per parasite line. Mean values are highlighted in red. Statistical evaluation used unpaired two-tailed students t-test (n.s., not significant). Representative Giemsa-stained images of WT and KO parasites are shown on the right. F, G) Mitochondrial morphology as visualized by MitoTracker Red staining (red) in WT and PNPLA2-KO parasites. In (F) mitochondrial morphology was evaluated between 24 and 40 hpi and parasites were divided according to their number of nuclei. Shown are mean values of two independent experiments. The total number of schizonts analyzed in each category is shown at the bottom of each graph. Representative images of the different mitochondrial morphologies observed are displayed on the left. In (G) WT and KO schizonts were arrested from 40 to 48 hpi with C2 to prevent egress and the percentage of schizonts having at least one abnormal mitochondrial accumulation determined. Shown are means +/- SD of four independent experiments, in which a total of 381 WT and 376 KO schizonts were analyzed. Statistical evaluation used unpaired students t-test (**** p < 0.0001). Representative images of WT and KO schizonts are shown on the left. DAPI-stained nuclei are shown in blue. A typical mitochondrial accumulation observed in KO parasites is arrowed. All scale bars, 5 µm.

We next characterized the growth phenotype of PNPLA2-null parasites in more detail, examining the development of tightly synchronized intracellular parasites over the course of the erythrocytic cycle. This showed that PNPLA2-null parasites exhibited delayed development in comparison to WT parasites, as evident by microscopic quantification of Giemsa-stained thin blood films (Figure 4D). However, this also revealed that in those PNPLA2-null schizonts that developed to maturity, there was no significant decrease in daughter merozoite numbers as compared to WT schizonts (Figure 4E), indicating that loss of PNPLA2 leads to delayed but not compromised parasite maturation.

Given the mitochondrial localization of PNPLA2, we next studied mitochondrial development in the PNPLA2-null parasites. Microscopic examination of mitochondria during trophozoite and schizont development using MitoTracker Red revealed mitochondrial abnormalities in the mutant parasites in the form of accumulations that first became evident following the first round of nuclear division (2 nuclei) and were further pronounced at later stages of schizogony (3 and more nuclei) (Figure 4F). To further analyze and quantify this phenotype, we performed an end-point analysis by arresting egress of WT and PNPLA2-null schizonts for 8 hours using C2 and then quantifying mitochondrial morphology using MitoTracker Red staining. This showed that whilst most segmented WT schizonts displayed the typical comma-like structure of divided mitochondria, ∼80% of PNPLA2-null schizonts showed abnormal mitochondrial accumulations, together indicating that PNPLA2 is involved in mitochondrial morphogenesis (Figure 4G).

### Confirmation of the PNPLA2-null phenotype by conditional gene disruption

To further analyze the function of PNPLA2, and to establish whether the observed defects were detectable immediately following gene disruption, we next targeted the *pnpla2* gene using the DiCre-based conditional KO approach. To this aim, we again used Cas9-assisted double homologous recombination to flox the sequence encoding the C-terminal half of the PNPLA2 coding sequence (harboring the catalytic PNPLA domain), simultaneously appending a 3xHA epitope tag to the gene (Figure 5A). As previously, this manipulation was performed in the DiCre-expressing B11 *P. falciparum* line. Two clonal transgenic parasite lines (called PNPLA2:HA:loxPint clones C9 and D11) were isolated and the expected genomic modifications confirmed by PCR (Figure 5 – figure supplement 1).

**Figure 5.**
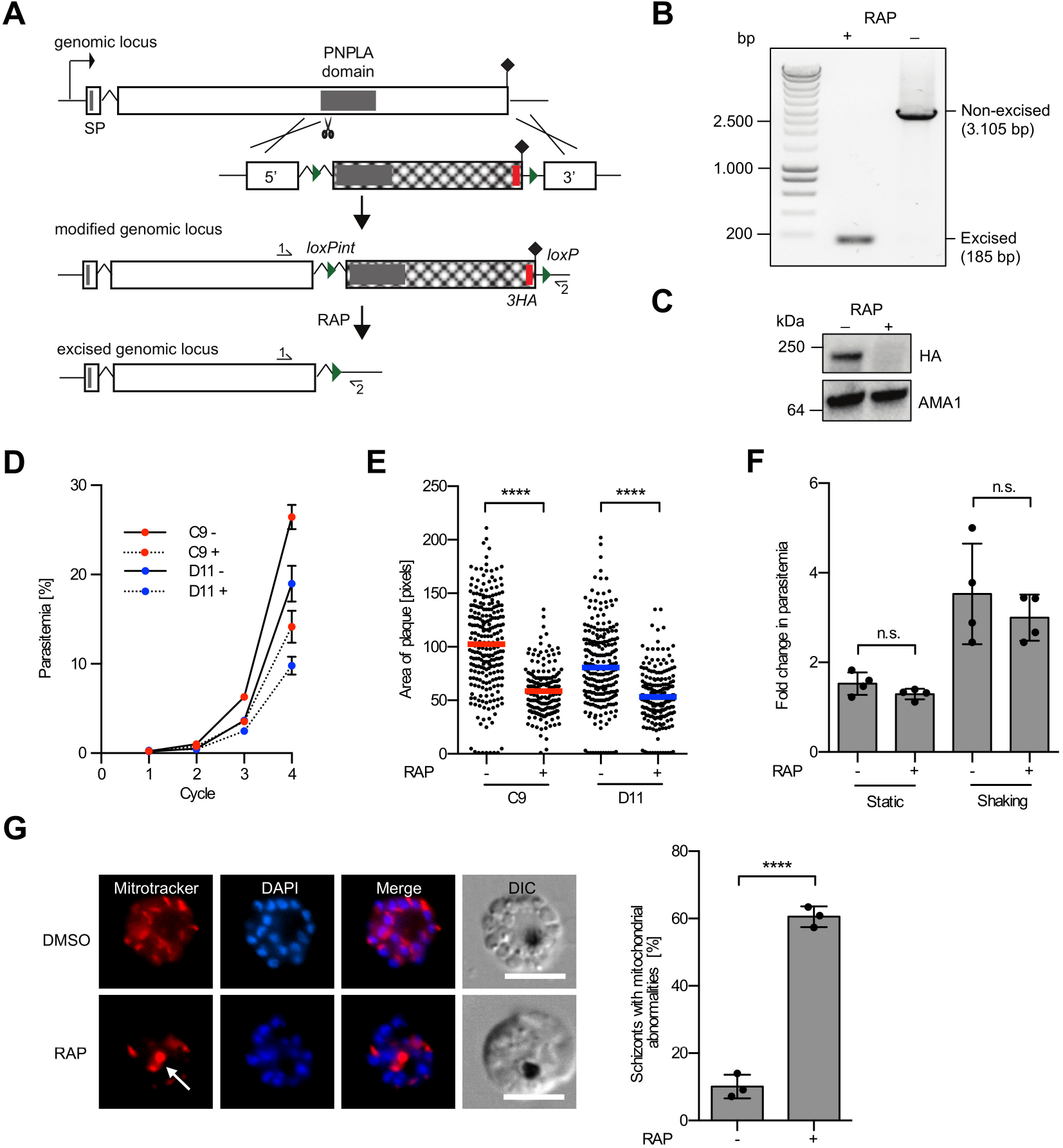
Conditional gene disruption confirms a key role of PNPLA2 in parasite growth and mitochondrial morphogenesis. A) Schematic of the strategy used to make a conditional PNPLA2-KO line (PNPLA2:HA:loxPint). The PNPLA domain (dark grey) was floxed by introducing a *loxP*-containing intron (loxPint) upstream of the domain and a second *loxP* site downstream of the translational stop site (lollipop). Sites of targeted Cas9-mediated double-stranded DNA break (scissors), left and right homology arms for homology-directed repair (5’ and 3’), introduced *loxP* sites (arrow heads), recodonized sequences (hatched) and 3xHA epitope (red) are indicated. RAP-induced DiCre-mediated excision results in removal of the functional domains. Primers 1 and 2 (half arrows) were used for diagnostic PCR. B) Confirmation of efficient gene excision by PCR. Samples were taken at 12 h post RAP or mock (DMSO) treatment of ring stage parasites. Expected PCR product sizes for non-excised and excised parasites are shown. Displayed are results of one representative experiment (out of five independent experiments). C) Western blot of C2-arrested mature schizonts (48 hpi) showing successful RAP-induced ablation of PNPLA2-HA expression in the erythrocytic cycle of RAP addition. AMA1 served as a loading control. Shown is one representative out of two independent experiments. D) Replication of mock- (solid line) and RAP-treated (dashed line) parasites from two clonal lines of PNPLA2:HA:loxPint parasites over four erythrocytic cycles. Shown are means +/- SD of three biological replicates. E) Replication of individual mock- and RAP-treated parasites from two clonal lines over five erythrocytic cycles measured as area of clonal plaques formed after 10 days of growth. Statistical evaluation used an unpaired students t-test (**** = P < 0.0001). F) Fold change in parasitemia after 4 h invasion of mock- and RAP-treated schizonts under static and shaking conditions. Shown are means +/- SD of two independent experiments with two biological replicates each. Statistical evaluation used unpaired students t-test (n.s., not significant). G) Mitochondrial morphology as visualized by MitoTracker Red (red) staining in mock- and RAP-treated PNPLA2:HA:loxPint parasites. Schizonts were arrested from 46 to 49 hpi with C2 to prevent egress and the percentage of schizonts having at least one abnormal mitochondrial accumulation was determined. Shown are means +/- SD of three independent experiments, in which a total of 481 mock- and 522 RAP-treated schizonts were analyzed. Unpaired students t-test was used for statistical evaluation (**** p < 0.0001). Representative images of mock- and RAP-treated schizonts are shown on the left. DAPI-stained nuclei, blue. The arrow indicates a typical mitochondrial accumulation observed in KO parasites. DIC, differential interference contrast. All scale bars, 5 µm.

RAP treatment of synchronous PNPLA2:HA:loxPint ring stage parasites resulted in the expected truncation of the PNPLA2 gene within a single erythrocytic cycle, as confirmed by PCR (Figure 5B) and western blot analysis (Figure 5C). Flow cytometry-based growth analysis revealed a reduction in replication rate of ∼50% in RAP-treated parasites over four erythrocytic cycles, as compared to mock-treated PNPLA2:HA:loxPint parasites (Figure 5D). To assess the longer-term viability of PNPLA2-null parasites, we used a plaque assay (Thomas et al., 2016), which provides a measure of replication over a period of ∼5 erythrocytic cycles by visualization of localized zones of RBC lysis in static parasite cultures. At limiting dilution parasitemia levels, each plaque arises from clonal expansion of a single parasite and the area of the plaque is proportional to the clonal replication rate. In both clonal lines C9 and D11, RAP treatment resulted in a 25-30% reduction in numbers of plaques formed (number of plaques: C9-RAP+, 180; C9-RAP-, 239; D11-RAP+, 633; D11-RAP-, 999). Additionally, we observed a 34-40% reduction in the average area of each plaque in the RAP-treated cultures (Figure 5E). Together with the results of the flow cytometry based growth assays, these results further highlight the importance of PNPLA2 for parasite proliferation. For more in-depth characterization, the clonal line D11 was used in all subsequent experiments.

To establish whether the reduced replication rate in PNPLA2-null parasites was due to inefficient egress or invasion, we isolated schizonts from RAP- and mock-treated PNPLA2:HA:loxPint cultures at the end of the cycle of treatment and incubated them with fresh RBCs under both static and shaking conditions. This showed no significant differences between the resulting increases in parasitemia (Figure 5F), suggesting that loss of PNPLA2 does not impair egress or invasion.

To test whether conditional disruption of PNPLA2 led to a mitochondrial development defect similar to that observed following direct gene disruption (Figure 4), we assessed mitochondrial morphology in mock- and RAP-treated PNPLA2:HA:loxPint schizonts by MitoTracker Red staining. This also revealed abnormal mitochondrial morphology in the RAP-treated schizonts (Figure 5G), confirming the importance of PNPLA2 in mitochondrial morphogenesis.

### Disruption of PNPLA2 impairs mitochondrial electron transport chain function

Given the mitochondrial localization of PNPLA2 and the KO-associated mitochondrial morphogenesis phenotype, we next aimed to characterize the susceptibility of our SLI-based PNPLA2-KO parasites towards several drugs that target mitochondrial functions to test whether disruption of PNPLA2 might impact the efficiency of these compounds. For this we performed a 96-hour SYBR-gold growth assay starting with trophozoite stage parasites and tested the growth of WT and PNPLA2-KO parasites under varying concentrations of drugs. Interestingly, PNPLA2-KO parasites showed decreased IC_50_ values for proguanil (13 fold) and the mitochondrial electron transport chain (mtETC) inhibitors atovaquone (5 fold), myxothiazol (6 fold) and antimycin A (7 fold) in comparison to WT parasites (Figure 6A-D). No increased sensitivity of KO parasites was seen for other drugs such as DSM1, dihydroartemisinin (DHA) and primaquine (Figure 6E-G), excluding a general increased drug susceptibility of the PNPLA2-KO parasites. Collectively, these data strongly suggest that disruption of PNPLA2 sensitizes parasites to antimalarial drugs that inhibit mitochondrial function. One main role of the malarial mtETC is to recycle ubiquinone that is necessary for ubiquinone-dependent enzymes, including dihydroorotate dehydrogenase (DHODH), which is the target of DSM1 and essential for the pyrimidine biosynthesis pathway (Goodman et al., 2017; Phillips et al., 2008). To test whether the ubiquinone pool was affected by disruption of PNPLA2, we treated parasites with increasing concentrations of the ubiquinone analog decylubiquinone (DCUQ). As reported previously (Ke et al., 2011), this treatment rescued an atovaquone-induced growth arrest of WT parasites in a dose-dependent manner. However, no rescue of PNPLA2 KO-parasite growth was observed upon DCUQ treatment (Figure 6H). This, together with the comparable susceptibility of PNPLA2-KO and WT parasites towards DSM1 suggests that disruption of PNPLA2 likely does not impair ubiquinone recycling.

**Figure 6.**
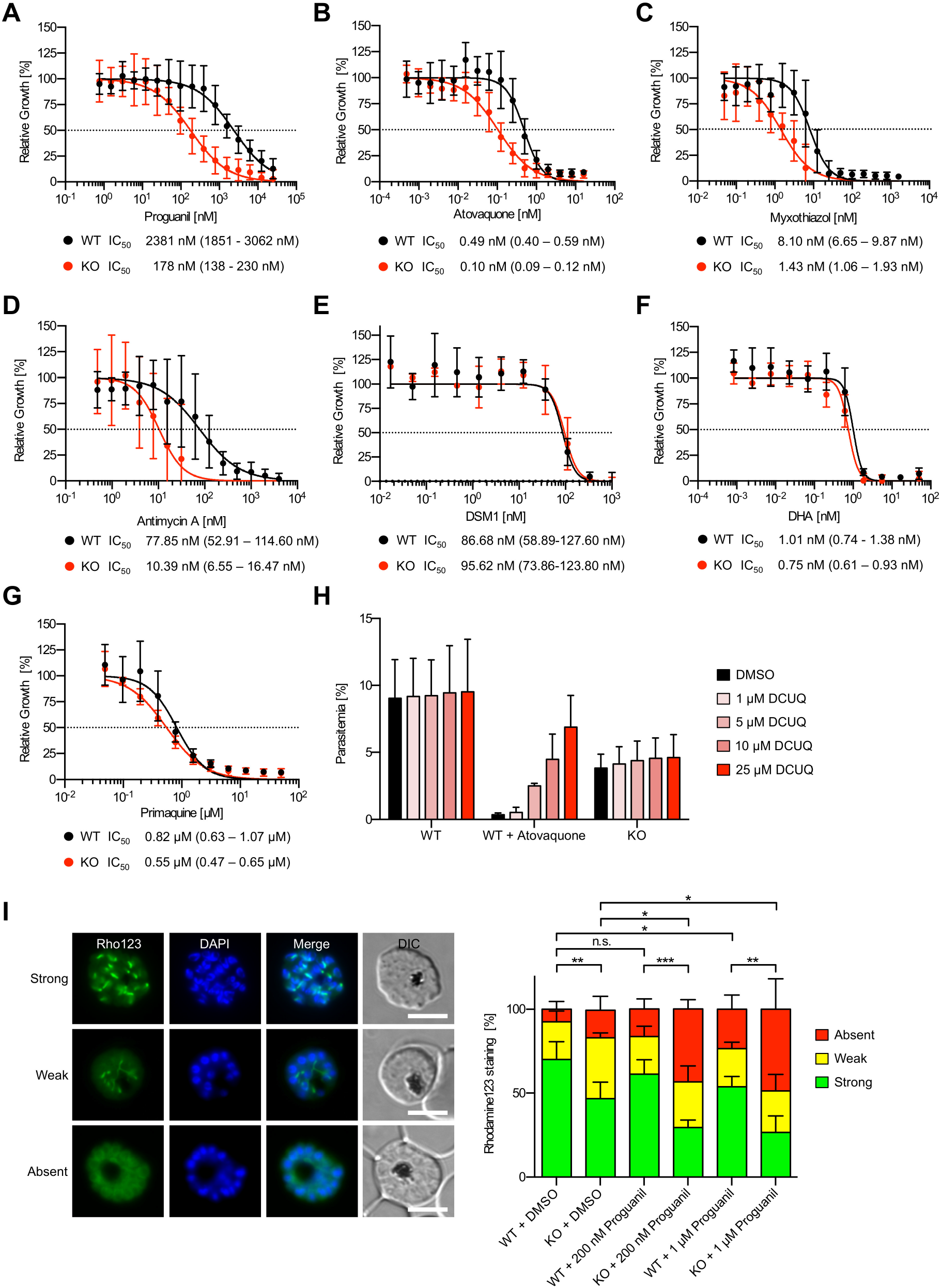
PNPLA2-KO parasites have a defect in the mtETC. A-G) Drug susceptibility assays of WT and PNPLA2-KO parasites using proguanil (A), atovaquone (B), myxothiazol (C), antimycin A (D), DSM1 (E), dihydroartemisinin (DHA, F), primaquine (G). Parasite growth was assessed by measuring DNA content using SYBR gold when exposed to varying concentrations of drugs for 96 h. The growth of DMSO-treated control parasites was set to 100%. Shown are means +/- SD of 3 to 6 independent experiments performed in duplicate. Calculated IC_50_ values with 95% confidence intervals are shown below each graph. H) The artificial electron acceptor decylubiquinone (DCUQ) does not rescue growth of PNLA2-KO parasites. WT and PNPLA2-KO parasites were grown in presence of various concentrations of DCUQ for two parasites cycles and parasitemia was evaluated using flow cytometry. As positive control, WT parasites were additionally treated with 1,15 nM atovaquone. Shown are means +/- SD of three independent experiments. I) PNPLA2-KO parasites have a defect in sustaining normal ΔΨm. C2-arrested WT and PNPLA2-KO schizonts that had been treated with DMSO (solvent control), 200 nM or 1 µM of proguanil were stained with the mitochondrial potentiometric dye rhodamine123 (Rho123, green) and parasites with a strong, weak or absent mitochondrial rhodamine123 signal were quantified by fluorescence microscopy. Shown are means +/- SD of four independent experiments, in which a total of 352 to 414 schizonts were analyzed per cell line and condition. For statistical evaluation a one-way ANOVA followed by a Holm-Sidak multiple comparison test was performed (*p < 0.05; **p < 0.01; ***p < 0,001; n.s., not significant). Representative images are shown on the left. DAPI-stained nuclei are shown in blue. DIC, differential interference contrast. Scale bars, 5 µm.

Another important function for the mtETC in blood stage malaria parasites is to pump protons across the mitochondrial inner membrane to build up a proton electrochemical gradient, with the mitochondrial membrane potential, ΔΨm, as its major component. The energy saved in ΔΨm is then used by the mitochondrion to import proteins and metabolic precursors from the cytosol, to maintain critical biochemical pathways such as the generation of iron-sulfur clusters (Painter et al., 2007; Vaidya and Mather, 2009). To test whether the ΔΨm is affected in PNPLA2-KO parasites, we incubated WT and PNPLA2-KO schizonts with the mitochondrial potentiometric dye rhodamine123 as previously described (Matz et al., 2018) and analyzed them by fluorescence microscopy. This allowed differentiation of parasites with i) strong mitochondrial rhodamine123 signal, ii) visible but weak mitochondrial staining and iii) absent fluorescence or cytoplasmic/peripheral signal. This analysis revealed significantly more PNPLA2-KO parasites showing an abnormal (weak or absent) rhodamine123 signal in comparison to WT parasites, indicating that disruption of PNPLA2 impairs a mitochondrial function that is necessary to sustain normal ΔΨm (Figure 6I). Given that, of the drugs tested, PNPLA2-KO parasites showed the highest level of hypersensitivity towards proguanil (Figure 6A), parasites were also treated with increasing concentrations of this drug to test how this affected ΔΨm. In line with our drug susceptibility data, treatment of PNPLA2-KO parasites with 200 nM proguanil led to a statistically significant increase in parasites showing an abnormal rhodamine123 staining in comparison to DMSO-treated PNPLA2-KO parasites, while the differences observed for WT parasites at this concentration of the drug did not reach statistical significance. Of note, treating WT parasites with 1 µM proguanil also significantly increased the percentage of parasites showing an abnormal rhodamine123 signal, suggesting that proguanil treatment alone does impact on ΔΨm in WT parasites. This stands in contrast to a previous study performed in a rodent malaria model that found that proguanil alone does not influence ΔΨm (Srivastava and Vaidya, 1999). Collectively, our data imply that PNPLA2-KO parasites may have a defect in the mETC that leads to an impaired ability to sustain ΔΨm and thereby renders parasite hypersensitive to mETC inhibitors.

## DISCUSSION

Phospholipases are ubiquitously occurring enzymes that catalyze the cleavage of phospholipid molecules. As a result, these enzymes are involved in diverse physiological processes including remodeling of cellular membranes, lipid-mediated signal transduction processes, cell proliferation and virulence (reviewed in (Flammersfeld et al., 2017)).

The focus of this study was a systematic functional characterization of 20 putative *P. falciparum* phospholipase genes for which mass-spectrometry data unambiguously indicated expression during asexual blood stage development. We first used a SLI-based gene disruption strategy to show that out of the 20 genes, 15 could be readily disrupted without loss of parasite viability, pointing to a high level of redundancy within this class of enzymes in *Plasmodium*. For the five remaining genes no transgenic knockout parasites were obtained, suggesting a possible critical role in parasite growth. Of these five genes, two previous studies concluded that the PNPLA1 (PF3D7_0209100) was dispensable for asexual parasite growth (Flammersfeld et al., 2019; Singh et al., 2019). Our essentiality data are in good agreement with a recent genome-wide saturation mutagenesis screen in *P. falciparum* (Zhang et al., 2018), except that we readily obtained gene disruption of five putative phospholipases that were predicted to be essential by the genome-wide KO screen.

Functional redundancy between phospholipases is well established, including an excellent example in *Listeria*, where individual disruption of two phospholipases resulted in moderate effects on infectivity to mice (2–20 fold reduction), whilst simultaneous disruption of both phospholipases in combination severely impaired infectivity (500 fold reduction) (Smith et al., 1995). Revealing the interplay between putative phospholipases in the malaria parasite by generation of double or even triple null parasites will be an interesting approach to discovering functional interplay within this enzyme family.

Some of the non-essential putative phospholipases identified in our work have been previously studied. The homologue of PF3D7_0629300 in the rodent malaria model *P. berghei* (*Pb*PL, PBANKA_1128100) exhibits phospholipase and membranolytic activity *in vitro* and has been implicated in cell traversal by sporozoites and disruption of the liver stage PVM during parasite egress from hepatocytes (Bhanot et al., 2005; Burda et al., 2015). The sphingomyelin phosphodiesterase (PF3D7_1238600) was identified as a PLC able to hydrolyze sphingomyelin and lysocholinephospholipids, and inhibitor studies using scyphostatin were used to argue for an essential role of the enzyme during asexual growth (Hanada et al., 2002). Our study now provides unambiguous reverse genetic evidence that PF3D7_1238600 is dispensable for parasite proliferation, suggesting that scyphostatin has additional targets within the parasite. PF3D7_0709700, previously designated as prodrug activation and resistance esterase *Pf*PARE, was shown to have esterase activity to activate esterified pepstatin, a peptidyl inhibitor of malarial aspartyl proteases (Istvan et al., 2017). *Pf*PARE active site mutants were not impaired in asexual blood stage growth (Istvan et al., 2017), mirroring our successful gene disruption. Finally, the two non-essential putative lysophospholipases PF3D7_1001400 and PF3D7_1001600 both contain a PEXEL motif and were shown to be exported into the host RBC, although their molecular functions have not yet been determined (Spillman et al., 2016).

Several previous studies have suggested that PI-PLC is essential for parasite blood stage proliferation, but definitive genetic evidence for this has been lacking. Earlier work has shown that the PI-PLC homolog in *P. berghei* (PBANKA_1211900) is refractory to genetic deletion (Raabe et al., 2011a). PI-PLCs are phosphodiesterases that participate in phosphatidylinositol 4,5-bisphosphate (PIP_2_) metabolism and lipid signaling pathways in a Ca^2+^-dependent manner (reviewed in (Kadamur and Ross, 2013)). Studies in malaria parasites have suggested that PI-PLC activity is involved in multiple processes ranging from gametocyte development and sporozoite motility to egress and invasion of merozoites by regulating Ca^2+^ release (Agarwal et al., 2013; Carey et al., 2014; Raabe et al., 2011b; Singh et al., 2010). PI-PLC was shown to likely act downstream of the parasite PKG, which regulates egress and activity of which promoted hydrolysis of the PI-PLC substrate PIP_2_ (Brochet et al., 2014). However, all of these studies relied on the use of the small compound inhibitor U73122, for which the degree of selectivity for PI-PLC is unclear, given that in numerous other systems the compound has the potential to modulate Ca^2+^ homeostasis independently of PI-PLC inhibition (Macmillan and McCarron, 2010; De Moel et al., 1995; Mogami et al., 1997).

Using two distinct conditional gene targeting approaches we now provide unambiguous genetic evidence that PI-PLC is essential for *P. falciparum* asexual blood stage proliferation. Both conditional inactivation techniques resulted in a defect in trophozoite to schizont conversion, as well as impaired development of schizonts. In line with this maturation phenotype, our lipidomic analysis showed a significantly reduced lipid content in the PI-PLC deficient parasites, while only a slight impact on the overall lipid profile was visible. It might be interesting, although experimentally challenging, to directly probe the enzymatic activity of PI-PLC by quantitation of PIP_2_ and inositol 1,4,5-triphosphate (IP_3_) using our PI-PLC deficient parasites.

The maturation phenotype of our PI-PLC deficient parasites is reminiscent of that seen in the related apicomplexan parasite *Toxoplasma gondii*, where conditional ablation of *Tg*PI-PLC caused significant morphological abnormalities during lytic stage growth (Bullen et al., 2016). Together, these findings therefore support a function for PI-PLC in a yet incompletely understood role that is necessary for daughter cell formation in these two apicomplexan genera and that occurs long before the role of PI-PLC in egress and invasion.

Of the 15 non-essential putative phospholipases analyzed in this study, only disruption of PNPLA2 (PF3D7_1358000) led to a growth defect. PNPLAs are highly conserved enzymes of prokaryotic and eukaryotic organisms with a broad physiological role (reviewed in (Wilson and Knoll, 2018)). Apart from PNPLA2, the *P. falciparum* genome encodes three other putative PNPLA enzymes (PF3D7_0209100, PF3D7_0218600, PF3D7_0924000), all of which appear to be redundant for asexual blood stage multiplication (Figure 1, (Flammersfeld et al., 2019; Singh et al., 2019)). Interestingly, PNPLA1 (PF3D7_0209100) seems to be critical in gametocyte induction (Flammersfeld et al., 2019) and gametogenesis (Singh et al., 2019).

Using GFP-tagging of the endogenous gene product, we unequivocally showed that PNPLA2 localizes to the parasite mitochondrion, in line with the fact that the N-terminus of PNPLA2 contains a predicted mitochondrial targeting signature sequence (Claros and Vincens, 1996). Interestingly, the putative *T. gondii* orthologue of PNPLA2 (TGME49_231370) localizes to the apicoplast and its conditional knockdown led to a rapid apicoplast loss due to deregulated lipid homeostasis in this organelle (Lévêque et al., 2017). Here we demonstrate that disruption of *P. falciparum* PNPLA2 impairs blood stage proliferation and that PNPLA2 deficient parasites display mitochondrial abnormalities, indicating a putative function of PNPLA2 in mitochondrial morphogenesis.

We further show that PNPLA2-KO parasites are hypersensitive to drugs that target the mtETC and that they have a defect in sustaining ΔΨm, which both together argues for a defect somewhere in the mtETC. Although the molecular dissection of this defect requires further investigation, it is likely that it occurs downstream of complex III, for instance in the electron transport from complex III to complex IV or in the activity of the latter. This assessment is based on the facts that i) we did not observe hypersensitivity towards DSM1 and ii) we could not rescue the KO associated growth defect by treatment with the ubiquinone analog DCUQ. In this regard it is interesting to note that certain phospholipids of the inner mitochondrial membrane, in particular cardiolipin and phosphatidylethanolamine, are important for full activity of the mtETC and the efficient generation of ΔΨm by affecting supercomplex-formation between complex III and IV and by regulating complex IV activity (Böttinger et al., 2012; Pfeiffer et al., 2003). One plausible explanation for the defects observed in PNPLA2-KO parasites would thus be that KO parasites have a deregulated mitochondrial phospholipid metabolism, which negatively affects the activity of the mtETC. Although this still needs to be shown, a role of PNPLA2 in regulation of mitochondrial phospholipids is likely, especially given the fact that the putative orthologue of PNPLA2 in *T. gondii* is implicated in lipid homeostasis (Lévêque et al., 2017). In further support of this hypothesis, a similar function was previously described for the calcium-independent phospholipase A2γ in mice that also localizes to the mitochondrion and disruption of which was associated with deregulated cardiolipin metabolism and reduced complex IV activity (Mancuso et al., 2007). Interestingly, reduced complex IV activity in *Caenorhabditis elegans* and mammalian cells has been shown to trigger mitochondrial hyperfusion to transiently compensate for a decreased activity of the mtETC (Rolland et al., 2013). A similar mechanism might also be induced in the PNPLA2-deficient parasites, perhaps explaining the mitochondrial accumulations that we consistently observed in these parasites. Alternatively, these accumulations might be the result of a deregulated lipid metabolism per se, since mitochondrial morphology is regulated by the relative rates of mitochondrial fusion and division, processes that are performed by a conserved protein machinery and that are both highly dependent on the mitochondrial phospholipid composition (reviewed in (Furt and Moreau, 2009)).

In addition to mtETC inhibitors, PNPLA2-KO parasites also became hypersensitive towards proguanil, which is combined together with atovaquone in the registered antimalarial formulation Malarone. Although proguanil’s mechanism of action is not completely understood, combination studies have shown that proguanil acts by lowering the concentration at which atovaquone collapses parasite ΔΨm (Painter et al., 2007; Srivastava and Vaidya, 1999). It is hypothesized that this process is connected to an ATP synthase function, which only becomes essential when the mtETC is inhibited. During mtETC inhibition, ATP synthase could maintain ΔΨm by operating in reverse and it may be this function that is inhibited by proguanil (Painter et al., 2007). Given the high level of hypersensitivity of PNPLA2-deficient parasites towards proguanil, it is tempting to speculate that the proguanil-sensitive pathway for maintaining ΔΨm in presence of mtETC inhibition might partially compensate for the PNPLA2-KO-associated defect in the mtETC and that this is one reason why PNPLA2-KO parasites are still viable.

In conclusion, our study provides a functional framework of phospholipases in the clinically relevant blood stages of *P. falciparum*. Our work identifies PI-PLC as an essential regulator of parasite maturation, and demonstrates a critical role for a malarial PNPLA in mitochondrial morphogenesis and function. Collectively, our findings expand the repertoire of functions that phospholipases may perform in this important human pathogen.

## MATERIALS AND METHODS

### Cloning of SLI plasmids

For generation of SLI-based TGD constructs, 312-954 bp immediately downstream of the start ATG of the target genes were amplified by PCR, starting with a stop codon, to serve as homology regions for single-crossover based integration. PCR products were cloned using NotI/MluI into pSLI-TGD (Birnbaum et al., 2017) to generate the final targeting plasmids.

For generation of the PI-PLC knocksideways construct pSLI-PF3D7_1013500-KS, the C-terminal 985 bp of the *pi-plc* gene were amplified by PCR using primers PF3D7_1013500-tag-fw/PF3D7_1013500-tag-rev, starting with a stop codon, and cloned into pSLI-sandwich (Birnbaum et al., 2017) using NotI/AvrII.

For generation of the PNPLA2 GFP-tagging construct pSLI-PF3D7_1358000-GFP, the C-terminal 1,063 bp of the *pnpla2* gene were amplified by PCR using primers PF3D7_1358000-tag-fw/PF3D7_1358000-tag-rev and cloned into pSLI-TGD (Birnbaum et al., 2017) using NotI/MluI.

Phusion High-Fidelity DNA polymerase (New England BioLabs) was used for all plasmid constructions and all plasmid sequences were confirmed by Sanger sequencing. For primer sequences see Supplementary file 1.

### Cloning of plasmids for conditional gene KO

Gene segments containing the catalytic domains of the *pi-plc* and *pnpla2* genes were replaced by a synthetic, modified version using Cas9-enhanced homologous recombination by transfecting a guide plasmid and a linearized repair plasmid into the DiCre-expressing *P. falciparum* line B11 (Perrin et al., 2018). A 2,417 bp and 2,352 bp long gene segment were chosen in *pi-plc* and *PNPLA2* respectively.

Two single guide RNA (sgRNA) inserts per target gene were generated by annealing oligo pairs PF3D7_1013500_gRNA01.F/ PF3D7_1013500_gRNA01.R and PF3D7_1013500_gRNA02.F/ PF3D7_1013500_gRNA02.R for PI-PLC and PF3D7_1358000_gRNA01.F/ PF3D7_1358000_gRNA01.R and PF3D7_1358000_gRNA02.F/ PF3D7_1358000_gRNA02.R for PNPLA2, which were subsequently ligated into the BbsI-digested plasmid pDC2-Cas9-hDHFRyFCU plasmid (Knuepfer et al., 2017) which contains sequences encoding Cas9, single guide RNA (sgRNA) and the drug selectable marker hDHFR (human dihydrofolate reductase)/yFCU (yeast cytosine deaminase/uridyl phosphoribosyl transferase).

Repair plasmids were designed such that they had i) ∼500 bp native sequences on either side of the targeted gene segment to serve as homology arms, ii) a short synthetic intron containing a *loxP* site (loxPint) upstream of the targeted gene segment, iii) the recodonized version of the targeted gene segment with the PAM sites destroyed, iv) a triple-hemagglutinin (3HA) epitope tag just prior to the gene translational stop codon, and v) another *loxP* site following the translational stop codon. To target PI-PLC, the above designed construct was synthesized as two parts (2,866 bp and 791 bp) and combined by restriction-ligation (using HindIII and XhoI enzymes) to create pREP-piplc-3HA-loxPint. Similarly, to target PNPLA2, the designed construct was synthesized as two parts (2,421 bp and 1,109 bp) and inserted subsequently into a pCR-blunt vector (Thermo Fisher Scientific) using restriction-ligation with XhoI/ApaI and XhoI/PstI respectively. The synthesized construct did not contain the 3’ homology arm, which was therefore amplified from B11 genomic DNA (amplification primers: PF3D7_1358000_3hom_F and PF3D7_1358000_3hom_R) and added to the construct by restriction-ligation using NheI/ApaI sites to create pREP-PNPLA2-3HA-loxPint.

Synthetic gene constructs were synthesized by GeneArt (Thermo Fisher Scientific). Phusion High-Fidelity DNA polymerase (New England BioLabs) was used for all plasmid constructions and all plasmid sequences were confirmed by Sanger sequencing. For sequences of all primers and synthetic gene constructs see Supplementary file 1.

### P. falciparum culture

Blood stages of 3D7 *P. falciparum* parasites were cultured in human RBCs. Cultures were maintained at 37°C in an atmosphere of 90% nitrogen, 5% carbon dioxide and 5% oxygen (DiCre-based KO lines) or in an atmosphere of 94% nitrogen, 5% carbon dioxide and 1% oxygen (all other parasite lines) using RPMI complete medium containing 0.5% Albumax according to standard procedures (Trager and Jensen, 1976).

### Generation of SLI-based parasite lines

For transfection of constructs, Percoll (GE Healthcare)-enriched synchronized mature schizonts of 3D7 parasites were electroporated with 50 µg of plasmid DNA using a Lonza Nucleofector II device (Moon et al., 2013). Transfectants were selected in medium supplemented with 3 nM WR99210 (Jacobus Pharmaceuticals), 0.9 µM DSM1 (BEI Resources) or 2 µg/ml blasticidin S (Invitrogen). For generation of stable integrant cell lines, parasites containing the episomal plasmids selected with WR99210 were grown with 400 µg/ml Neomycin/G418 (Sigma) to select for integrants carrying the desired genomic modification as described previously (Birnbaum et al., 2017). Each WR-resistant parasite culture was routinely placed under neomycin selection in three independent experiments using three culture dishes each time and was followed up for 60 days to monitor the appearance of viable transgenic parasites (expected to represent parasites in which the targeted gene was disrupted). Successful integration was confirmed by diagnostic PCR using FIREpol DNA polymerase (Solis BioDyne). For primer sequences see Supplementary file 1.

### Generation of conditional KO parasite lines

All transgenic *P. falciparum* DiCre-based KO parasite lines used in this study were based on the DiCre-expressing *P. falciparum* clone B11, derived from the 3D7 parasite line (Perrin et al., 2018). Two transfections (one per guide RNA) were performed for each gene target. Mature schizonts enriched using Percoll (GE Healthcare) were electroporated with 20 ug of guide plasmid and 60 ug of linearized repair plasmid using an Amaxa 4D electroporator and P3 Primary cell 4D Nucleofector X Kit L (Lonza) using programme FP158 as described (Collins et al., 2013). 24 hours post-transfection, the culture medium was replaced with fresh medium containing WR99210 (2.5 nM), which was withdrawn after 4 days. Once drug-resistant parasites appeared (in about 2 weeks), they were cloned by limiting dilution using a plaque-based method (Thomas et al., 2016). Successful integration was confirmed by diagnostic PCR using GOtaq Hot Start Green Master Mix (Promega). For primer sequences see Supplementary file 1.

### Fluorescence microscopy

Mitochondria were stained by incubation of parasites in 20 nM MitoTracker Red CMXRos (Invitrogen) in culture medium for 15 min at 37°C. For staining of nuclei, parasites were incubated with 1 µg/ml DAPI (Sigma) in culture medium for 15 min at 37°C. DiCre-based conditional KO parasites were imaged using a Nikon Eclipse Ni-E widefield microscope, equipped with a Hamamatsu C11440 digital camera and a 100x/1.45NA oil immersion objective. All other parasite lines were imaged on a Leica D6B fluorescence microscope, equipped with a Leica DFC9000 GT camera and a Leica Plan Apochromat 100x/1.4 oil objective. Image processing was performed using ImageJ.

### Western blot

For western blot analysis, parasites were Percoll-enriched, washed and lysed with saponin. The resulting parasite pellets were solubilized in five volumes of a denaturing solubilization buffer (1% (w/w) SDS in 50 mM Tris-HCl, pH 8.0, 5 mM EDTA, 1 mM PMSF) with sonication. Samples were immediately boiled for 5 min, clarified by centrifugation at 12,000 × *g* for 20 min and subjected to SDS-PAGE. Proteins were transferred to nitrocellulose membranes. Membranes were then blocked in 3% BSA in PBS containing 0.2% Tween 20 before staining with rat anti-HA mAb 3F10 (Sigma, diluted 1:1,000) primary antibody in blocking buffer, then incubated with biotin-conjugated anti-rat antibody (Roche, diluted 1:8,000) in blocking buffer followed by horseradish peroxidase-conjugated streptavidin (Sigma, diluted 1:10,000). Antibody binding was detected using an Immobilon Western Chemiluminescent HRP Substrate (Millipore) and visualized using a ChemiDoc Imager (Bio-Rad) with Image Lab software (Bio-Rad). AMA1 and PKG were probed as loading controls using a rabbit anti-AMA1 antibody (Collins et al., 2009) (diluted 1:500) and a rabbit polyclonal human-PKG antibody (Enzo Lifesciences, diluted 1:1,000) respectively, followed by a HRP-conjugated goat anti-rabbit secondary antibody (Sigma, diluted 1:3,000).

### Immunofluorescence analysis

For IFA of PI-PLC-GFP-KS parasites, air dried thin blood films were fixed for 3 min in icecold methanol. After rehydration in PBS and blocking in 3% BSA/PBS, they were stained with mouse anti-AMA1 antibody (clone 1F9 (Coley et al., 2001), diluted 1:1,000) in blocking buffer, followed by staining with anti-mouse-AlexaFluor488 antibody (Invitrogen, 1:2,000) additionally containing 1 µg/ml DAPI in blocking buffer. Finally, DAKO mounting solution was added and slides were covered with a coverslip.

For IFA of PI-PLC:HA:loxPint parasites, air dried thin blood films were fixed with 4% paraformaldehyde in PBS for 30 min at RT, permeabilized with 0.1% (v/v) Triton X- 100 in PBS for 10 min, and blocked overnight in 4% BSA/PBS. Samples were probed with rat anti-HA 3F10 (Sigma, 1:500) in 4% BSA/PBS. Bound primary antibodies were detected using biotin-conjugated anti-rat antibody (Roche, 1:1,000) and AlexaFluor594-conjugated streptavidin (Life Technologies, 1:1,000) in 4% BSA/PBS. Slides were mounted in ProLong Gold Antifade Mountant with DAPI (Life Technologies).

### Analysis of SLI-based parasite lines

Schizont stage parasites of all analyzed parasite lines were isolated by Percoll enrichment and incubated with uninfected RBCs (5% hematocrit) for 3 h to allow rupture and invasion. Parasites were then treated with 5% sorbitol to remove residual unruptured schizonts, leading to a synchronous ring stage culture with a 3 h window. For growth analysis of PI-PLC-GFP-KS parasites, synchronous ring stage cultures were adjusted to ∼0.1% parasitemia and divided into two 2 ml dishes. To one of these dishes, rapalog (AP21967, Clontech) was added to a final concentration of 250 nM (rapalog was stored at −20°C as a 500 mM stock in ethanol, and working stocks were kept as 1:20 dilutions in RPMI at 4°C) while the other dish served as a control. Parasitemia was analyzed by flow cytometry at 1, 3, 5, and 7 days, when most of the parasites were at the trophozoite stage. After analysis on day 5, cultures were diluted 10-fold into fresh RBCs to prevent overgrowth. Medium with or without rapalog was changed daily.

For growth analysis of TGD-based KO lines, synchronous ring stage cultures were allowed to mature to trophozoites for one day. Parasitemia was then determined one day post-infection by flow cytometry and adjusted to exactly 0.1% starting parasitemia in a 2 ml dish. Medium was changed daily and growth of the parasite lines was assessed by flow cytometry after five days (two erythrocytic cycles). As a reference, WT 3D7 parasites were included in each assay.

For quantification of developmental stage and schizont analysis of PI-PLC-GFP-KS and PNPLA2-KO lines, synchronous ring stage cultures were diluted to ∼1-2% parasitemia in 2 ml dishes, which were either left untreated or treated with rapalog as described above in case of PI-PLC-GFP-KS parasites. Giemsa-stained blood films were prepared at 24, 40 and 48 hpi. For stage quantification, at least 20 fields of view were recorded using a 63x objective per sample. Erythrocyte numbers were then determined using the automated Parasitemia software (http://www.gburri.org/parasitemia/) and the number of the different parasite stages was manually counted on these images. For analysis of schizont morphology, cultures containing schizont stage parasites (40 hpi) were supplemented with the egress inhibitor compound 2 (1 µM; kindly provided by S. Osborne (LifeArc) and stored as a 10 mM stock in DMSO at −20°C). After 8 h, Giemsa-stained blood films were prepared and schizont morphology was investigated by light microscopy. For determining the number of merozoites per schizont, the cysteine protease inhibitor E64 (10 µM; Sigma) was added to schizonts at 40 hpi to prevent rupture of the host cell membrane. 6 to 8 h later, Giemsa smears were prepared and the number of merozoites per schizont was determined by light microscopy.

### Lipidomic analysis

Highly synchronous PI-PLC-GFP-KS ring stage parasite cultures were divided into four 10 ml plates. Two of these were treated with 250 nM Rapalog, while the other two were left untreated. Medium with or without Rapalog was replaced once per day. At 30 hpi and 40 hpi, parasitemia (3 – 7%) and the total number of erythrocytes were determined by flow cytometry for calculation of the absolute number of parasites per sample (50 – 140 x 10^6^ parasites). Per treatment and time point, parasites from one 10 ml dish were isolated by saponin lysis. Therefore, they were first washed in icecold PBS, followed by incubation in 0,03% saponin in PBS on ice for 10 min. After three washes in icecold PBS, parasite pellets were resuspended in 200 µl of PBS to which 800 µl of icecold LC-MS grade methanol (Merck) containing 0,1% (w/v) butylated hydroxytoluene (Sigma) were added. Samples were stored at -80°C until lipid extraction.

For the lipid extraction, samples were slowly thawed in ice cold water and sonified for 15 min. Directly afterwards aliquots were transferred into a new vial that correspond to 25 million parasites and the suspension was dried in a speed vac. To the dried cell pellets 50 µl water was added and rigorous stirred. The samples were then further homogenized using three freeze-thaw cycles consisting of a shock-freezing step in liquid nitrogen and 30 sec of sonification. Afterwards a mix of internal standards was added (Supplementary file 3). Lipid extraction was further performed according to earlier described lipid extraction method using MTBE (Matyash et al., 2008). Cholesterol was determined after acetylation as described (Liebisch et al., 2006). Shotgun lipidomics measurements were performed as described earlier (Eggers and Schwudke, 2018) using Q Exactive Plus (Thermo Fisher Scientific, Bremen, Germany) mass spectrometer coupled with the TriVersa NanoMate (Advion, Ithaca, USA). Lipid identification was performed with LipidXplorer 1.2.1 (Herzog et al., 2011) and post processing including quantitation was executed with lxPostman.

### Analysis of conditional KO parasite lines

Tightly synchronized ring stage cultures were divided into two dishes and treated with 100 nM rapamycin (Sigma, prepared as a 10 mM stock in DMSO) or DMSO only for 3 h at 37°C, following which the cultures received fresh medium. 24 h later, growth assays were set up for each treatment. For this, trophozoite stage parasites were assays, schizonts were isolated 48 h after the beginning of rapamycin treatment by Percoll enrichment and replicate cultures of each treatment were set up at ∼5% parasitemia with fresh RBCs. Parasites were allowed to invade for 4 h at 37°C under static or shaking (110 rpm) conditions. Giemsa smears were prepared at selected time points and parasite development and morphology assessed and quantified by light microscopy. In order to enrich the cultures with mature schizont stage parasites, parasites were treated at 46 hpi for 3 h with 1 µM C2 to arrest egress.

### Flow cytometry

For growth quantification of DiCre-based KO parasite lines, parasites were fixed with 0.1% glutaraldehyde/PBS and stained with SYBR Green I dye (1:10,000 dilution in PBS; Life Technologies) for 30 min at 37°C. Samples were analyzed in a BD Fortessa FACS instrument using the 530/30-blue detector configuration. Flow cytometry data was analyzed using FlowJo v10. Erythrocytes were gated based on their forward and side scatter parameters, and SYBR Green I stain-positive RBCs were identified using the 530/30-blue detector.

Flow cytometry-based analysis of growth of all other parasite lines was performed essentially as described previously (Malleret et al., 2011). In brief, 20 µl resuspended parasite culture was incubated with dihydroethidium (5 µg/ml, Cayman) and SYBR Green I dye (0.25 x dilution, Invitrogen) in a final volume of 100 µl medium for 20 min at RT protected from light. Samples were analyzed on a ACEA NovoCyte flow cytometer. RBCs were gated based on their forward and side scatter parameters. For every sample, 100,000 events were recorded and parasitemia was determined based on SYBR Green I fluorescence.

### Plaque assay

Long-term parasite replication rate as measured by plaque-forming ability was determined by diluting trophozoite stage cultures to a density of 10 parasites per well in complete medium with RBCs at a hematocrit of 0.75% as previously described (Thomas et al., 2016). This suspension was plated into flat bottom 96 well microplates (200 µl per well) and incubated under static conditions for 10 days in gassed humidified sealed modular chambers. Plaque formation was assessed by microscopic examination using a Nikon TMS inverted microscope (40x magnification) and documented using a Perfection V750 Pro scanner (Epson, Long Beach, CA). Plaques were counted by visual examination of the scanned images and plaque size quantified using the Lasso tool in Adobe Photoshop 2019.

### Transmission electron microscopy

PI-PLC:HA:loxPint parasites were treated at ring stage with RAP or DMSO, as described above, and allowed to develop to schizont stage. Schizonts were Percoll enriched and incubated with 1 µM C2 for 3 h. Samples were then fixed with 2.5% glutaraldehyde-4% formaldehyde in 0.1 M phosphate buffer (PB) for 30 min at RT. Schizonts were embedded in 3% low melting point agarose and the samples then cut into 1 mm^3^ blocks. These were then processed using a modified version of the NCMIR protocol (Deerinck et al., 2010). Briefly, blocks were washed in 0.1 M PB, post-fixed with 1% reduced osmium (1% OsO_4_/ 1.5% K_3_Fe(CN)_6_) for 1 h at 4^°^C, then washed in double distilled water (ddH_2_O). The blocks were incubated in 1% thiocarbohydrazide (TCH) for 20  min at RT, rinsed in ddH_2_O and further fixed with 2% osmium tetroxide for 30  min at RT. The blocks were then stained with 1% uranyl acetate at 4^°^C overnight, washed in ddH_2_O and stained with Walton’s lead aspartate for 30 min at 60^°^C. The blocks were washed in ddH_2_O and dehydrated stepwise using serial dilutions of ethanol: 30% and 50% at RT for 5 min each, then 70%, 90% and 2 x 100% for 10 min each. The blocks were infiltrated with a 4:1 mixture of propylene oxide (PO):Durcupan resin (Sigma) for 1 h at RT, followed by 1:1 and 1:4 mixtures for 1 h each at RT, then with 100% Durcupan resin for 48 h. Blocks were polymerised in fresh Durcupan resin at 60^°^C for 48 h. The samples were cut into 70 nm ultrathin sections using an ultramicrotome (UC7, Leica Microsystems UK) and picked up onto copper mesh grids (Agar Scientific). Images were obtained on a 120 kV transmission electron microscope (Tecnai G2 Spirit BioTwin, Thermo Fisher Scientific) using a charge-coupled device camera (Oneview, Gatan Inc.).

### Drug susceptibility assay

3D7-WT and SLI-based PNPLA2-KO parasites were synchronized to a 3 h time window as described for SLI-based parasite lines. At 24 hpi, parasitemia was determined by flow cytometry and drug susceptibility assays were set up in black 96-well microtiter plates (Thermo Scientific) with 0.1% starting parasitemia and 2% hematocrit in a final volume of 200 µl of medium. Hereby parasites were incubated with varying concentrations of the following drugs: proguanil (Sigma), atovaquone (Cayman), myxothiazol (Sigma), antimycin A (Sigma), DHA (Sigma), DSM1 (BEI Resources), primaquine (Cayman). Drugs were dissolved in PBS (primaquine, freshly prepared for every experiment) or DMSO (all other drugs) and then further diluted in culture medium. The final DMSO concentration did not exceed 0.25%. In each plate, infected RBCs in the absence of drugs and only treated with DMSO served as positive controls for parasite growth, whereas uninfected RBCs served as negative controls (background). After 96 h of incubation, inhibition of parasite growth was determined by measuring the fluorescence of SYBR Gold (invitrogen). Therefore, 100 µl/well supernatant were discarded without disturbing the RBC layer and 100 µl/well lysis buffer (20 mM Tris, 0.008% saponin, 0.08% Triton X-100, 1X SYBR Gold) were added. Plates were incubated in the dark for 2 h at RT before measuring fluorescence using the EnVision Multimode Plate Reader (PerkinElmer) with excitation and emission wavelengths of FITC 485 / FITC 535. IC_50_ values were calculated using nonlinear regression in GraphPad Prism (log(inhibitor) vs. normalized response – Variable slope).

### DCUQ assay

Growth of 3D7-WT and PNPLA2-KO parasites in presence of different concentrations of decylubiquinone (DCUQ, Cayman, stock prepared in DMSO) or DMSO was analyzed over two parasite cycles by flow cytometry as already described for the TGD-based KO parasite lines. As a positive control, WT parasites were treated with 1.15 nM atovaquone (IC_50_ value according to (Agarwal et al., 2017)) in addition to DCUQ/DMSO. Parasites were fed daily with fresh culture medium containing atovaquone, DCUQ or DMSO until analysis.

### Rhodamine123 based visualization of ΔΨm

Tightly synchronised ring stage cultures were treated with 200 nM proguanil, 1 µM proguanil or DMSO from 3 hpi until imaging. At 40 hpi, parasites were treated for 8 h with 1 µM C2 to prevent egress. Parasites were stained with rhodamine123 (Cayman) basically as previously described (Matz et al., 2018). In brief, parasites were incubated in 0.1 µg/ml rhodamine 123 and 1 µg/ml DAPI in culture medium for 30 min at 37°C. Afterwards, parasites were washed once in culture medium and further incubated at 37°C for another 30 min prior to live cell microscopy. The entire medium used during the staining procedure contained C2 and the respective amount of proguanil/DMSO. Image acquisition was performed using the same settings for all samples and at least 70 parasites per condition were imaged in each independent experiment.

### Statistical analysis

For statistical analysis of differences between two groups, paired or unpaired two-tailed students t-tests were used. For statistical analysis of differences between more than two groups, a one-way analysis of variance (ANOVA), followed by a Holm-Sidak multiple-comparison test was performed. All statistical tests were done in GraphPad Prism. P values of <0.05 were considered significant. Statistical details (n numbers, tests used, definition of the error bars) are described in the figure legends.

## DATA AVAILABILITY

All data generated or analyzed during this study are included in this published article and its supplemental material files. Detailed information on the lipidomics approach is available on the LIFS webportal (Schwudke, Shevchenko, Hoffmann, & Ahrends, 2017, https://lifs-tools.org). The preliminary LipidCompass accession number is LCE8.

## ACKNOWLEDGEMENTS

We thank Michael Foley for providing the monoclonal AMA1 antibody 1F9. The following reagent was obtained through BEI Resources, NIAID, NIH: DSM1, MRA-1161. We thank Anna Woitalla (RCB) for excellent technical support for the lipidomics analysis. We are grateful for expert support by Dr. Nils Hoffmann (Centrum for Biotechnology (CeBi/Tec), Universität Bielefeld) in data upload and data base management of the LIFS webportal. Images were acquired on microscopes of the CSSB imaging facility as well as the Electron Microscopy Science Technology Platform at the Francis Crick Institute. For the purpose of Open Access, the authors have applied a CC BY public copyright licence to any Author Accepted Manuscript version arising from this submission. PCB was funded by the Deutsche Forschungsgemeinschaft (DFG, German Research Foundation) – project number 414222880. AR was funded by a Marie Skłodowska Curie Individual Fellowship (Project number 751865). The work was also supported by funding to MJB from the Francis Crick Institute (https://www.crick.ac.uk/), which receives its core funding from Cancer Research UK (FC001043; https://www.cancerresearchuk.org), the UK Medical Research Council (FC001043; https://www.mrc.ac.uk/), and the Wellcome Trust (FC001043; https://wellcome.ac.uk/). The work was further supported by Wellcome ISSF2 funding to the London School of Hygiene & Tropical Medicine. Work in the DS lab was supported by LIFS2 project (FKZ 031L0108B) of the German Network for Informatic Infrastructure (de.NBI) and support by the German Center for Infection Research (TTU-TB, DZIF).

## AUTHOR CONTRIBUTIONS

Conceived and designed the experiments: PCB, AR, EP, MJB, TWG. Performed the experiments: PCB, AR, EP, SB, CS, LW, DS, AS, LMC. Analyzed the data: PCB, AR, EP, SB, CS, LW, JS, DS, MJB, TWG. Wrote the paper: PCB, AR, DS, MJB, TWG.

## FIGURES AND FIGURE SUPPLEMENTS

**Figure 1 - figure supplement 1.**
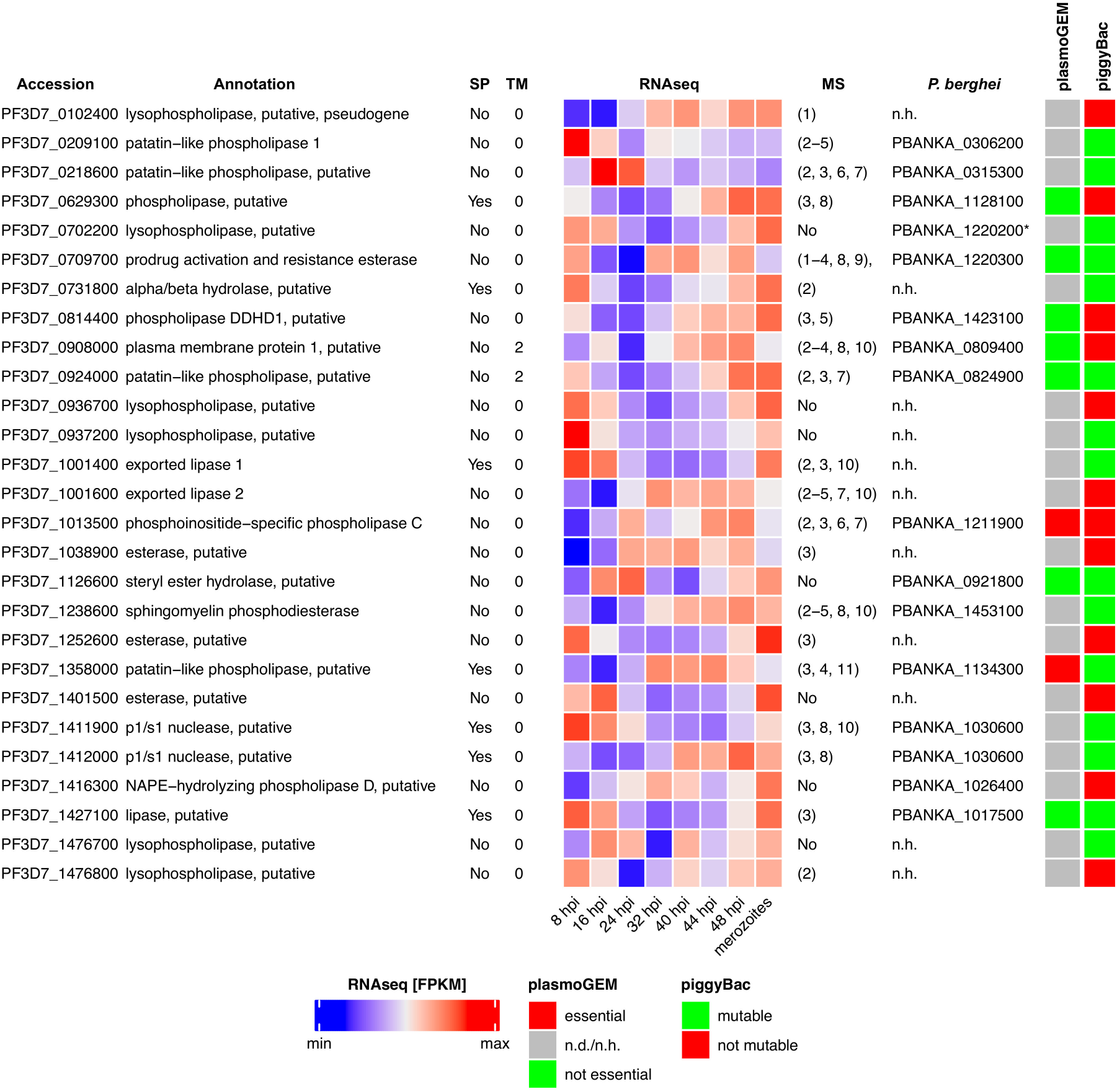
Putative phospholipases of *P. falciparum*. Mass-spectrometry (MS) expression data are based on the following references: 1) (Florens et al., 2002) 2) (Treeck et al., 2011) 3) (Pease et al., 2013) 4) (Bowyer et al., 2011) 5) (Solyakov et al., 2011) 6) (Lasonder et al., 2015) 7) (Lasonder et al., 2012) 8) (Oehring et al., 2012), 9) (Florens et al., 2004), 10) (Silvestrini et al., 2010), 11) (Cobbold et al., 2016). RNAseq expression data is derived from (Wichers et al., 2019). Orthologues in the rodent malaria model *P. berghei* were identified in PlasmoDB (Aurrecoechea et al., 2009) and are based on (Chen et al., 2006). Non-syntenic orthologues are marked with an asterisk. Results of the genome-wide KO screens in *P. berghei* (plasmoGEM) (Bushell et al., 2017) and *P. falciparum* using piggyBac-based mutagenesis (Zhang et al., 2018) are shown. SP, signal peptide; TM, transmembrane domain.

**Figure 1 – figure supplement 2.**
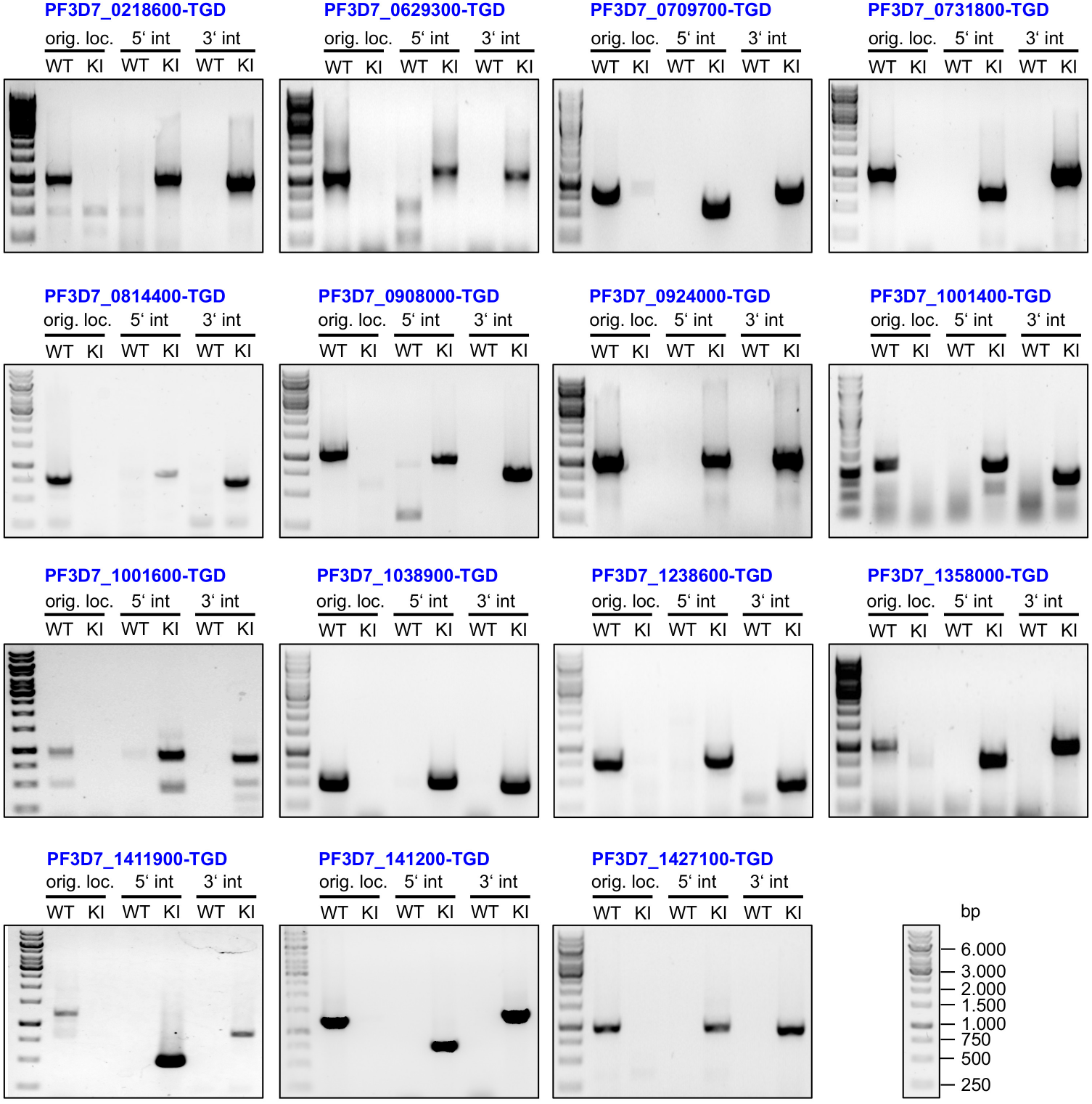
Integration PCRs of the TGD-based gene-deletion screen. Agarose gel electrophoresis of PCR products amplified from genomic DNA of the indicated parasite lines as well as unmodified WT parasites. Primers used are as indicated in Figure 1, demonstrating a product across the 5’ and 3’ integration junction (indicated as 5’ int and 3’ int, respectively) as well as quantitative absence of the original locus (‘orig. loc.’). Absence of this band indicates that no WT parasites remained in the parasite population. KI, knock in cell line. Fragment length of the markers is indicated once (bottom right). For primer sequences see Supplementary file 1.

**Figure 1 – figure supplement 3.**
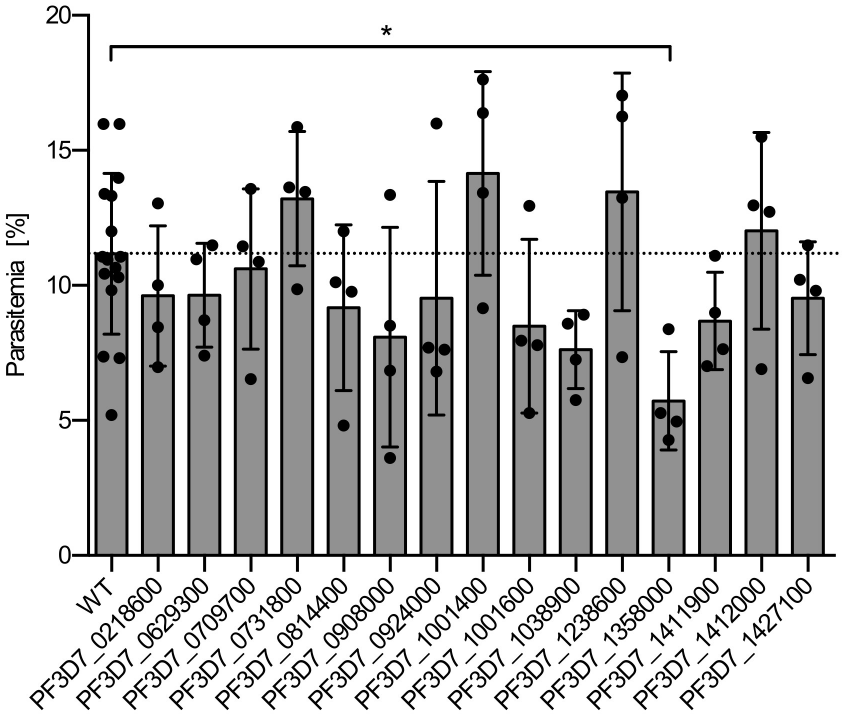
FACS-based growth analysis of synchronous phospholipase KO parasite lines after two erythrocytic cycles in comparison to 3D7 WT parasites. Raw parasitemia values with means +/- SD of four independent growth experiments per parasite line are shown. 3D7 WT parasites were included in each independent assay as a reference. For statistical analysis of growth rates of the different parasite lines in comparison to WT parasites, a one-way analysis of variance (ANOVA) followed by a Holm-Sidak multiple comparison test was performed. All statistically significant differences are indicated (* p < 0.05).

**Figure 2 – figure supplement 1.**
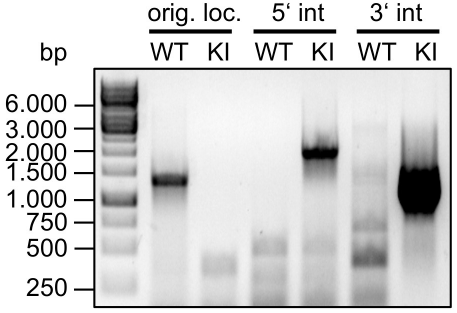
Integration PCR of PI-PLC-GFP-KS parasites. Agarose gel electrophoresis of PCR products amplified from genomic DNA of PI-PLC-GFP-KS as well as unmodified WT parasites. Primers used are as indicated in Figure 1, demonstrating a product across the 5’ and 3’ integration junction (indicated as 5’ int and 3’ int, respectively) as well as quantitative absence of the original locus (‘orig. loc.’). Absence of this band indicates that no WT parasites remained in the parasite population. KI, knock in cell line. For primer sequences see Supplementary file 1.

**Figure 2 – figure supplement 2.**
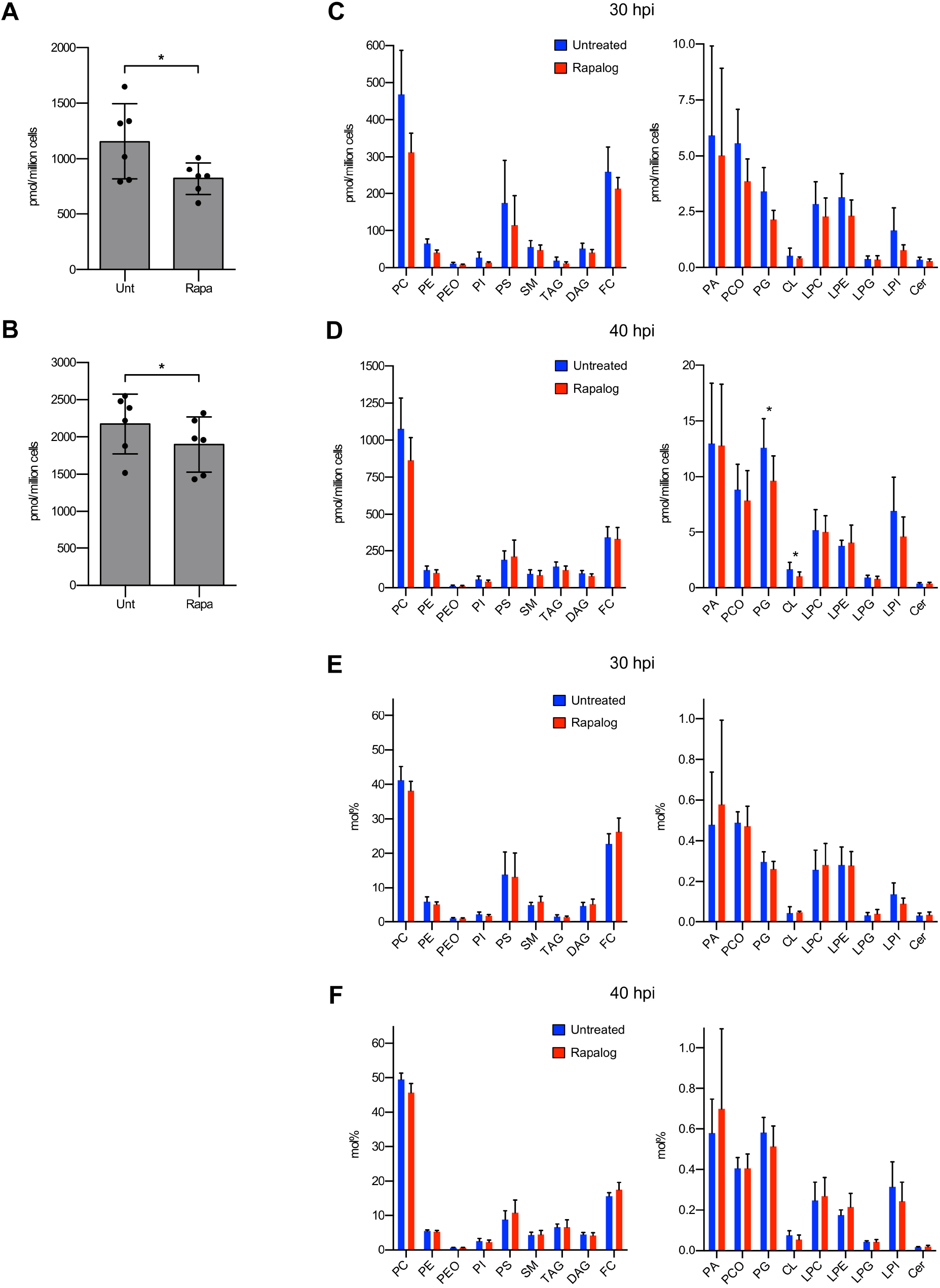
Lipidomic analysis of untreated and Rapa-treated PI-PLC-GFP-KS parasites. Parasites were grown in absence or presence of Rapa and harvested at 30 hpi and 40 hpi for lipidomic analysis. A, B) Total lipid amount of untreated and Rapa-treated PI-PLC-knocksideways parasites at 30 hpi (A) and 40 hpi (B). C, D) Absolute lipid levels of untreated and Rapa-treated PI-PLC-knocksideways parasites at 30 hpi (C) and 40 hpi (D). E, F) Relative abundance of lipid species in the total lipid composition (mol%) at 30 hpi (E) and 40 hpi (F). Data are based on 6 biological replicates. Means +/- SD are shown. Statistical evaluations were performed using paired two-tailed students t-test. For the data displayed in C-F, the Holm-Šídák method was used to correct for multiple comparisons. All statistically significant differences are indicated (* p < 0.05). PC, phosphatidylcholine; PE, phosphatidylethanolamine; PEO, alkyl-acylglycerophosphoethanolamines; PI, phosphatidylinositol; PS, phosphatidylserine; SM, sphingomyelin; TAG, triacylglycerol; DAG, diacylglycerol; FC, free cholesterol; PA, phosphatidic acid; PCO, alkyl-acylglycerophosphocholines; PG, phosphatidylglycerol; CL, cardiolipin, LPC, lysophosphatidylcholine; LPE, lysophosphatidylethanolamine; LPG, lysophosphatidylglycerol, LPI, lysophosphatidylinositol, Cer, ceramide. For a complete overview of all the results of the lipidomic analyses please see Supplementary file 2.

**Figure 3 – figure supplement 1.**
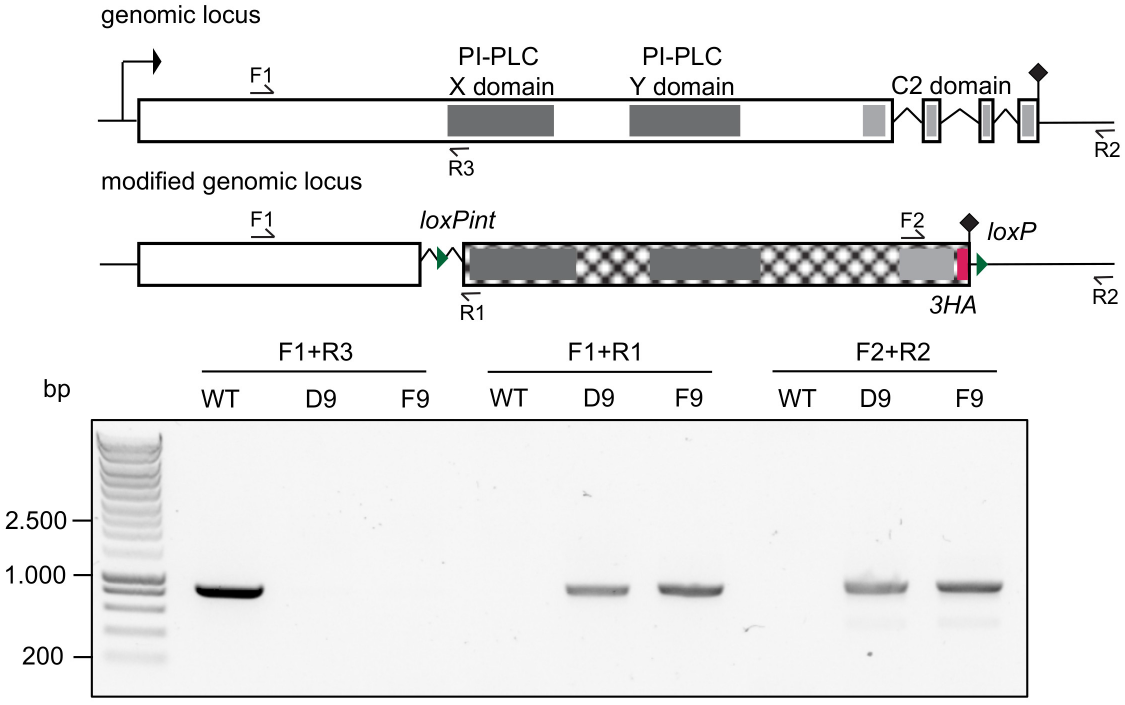
Integration PCR of PI-PLC:HA:loxPint parasites. Schematic of the *pi-plc* locus before and after CRISPR-Cas9-based gene editing is shown on top, while agarose gel electrophoresis of PCR products from unmodified WT and clonal modified parasite lines are displayed below. Primers used for confirming correct integration into the genome are indicated with arrows. For primer sequences see Supplementary file 1.

**Figure 4 – figure supplement 1.**
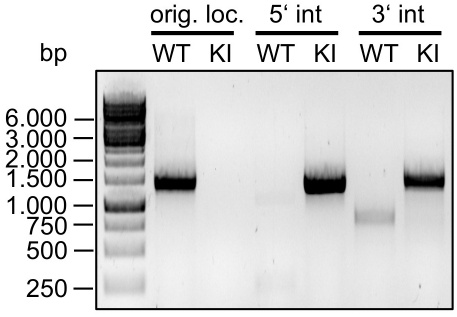
Integration PCR of PNPLA2-GFP parasites. Agarose gel electrophoresis of PCR products amplified from genomic DNA of PNPLA2-GFP as well as unmodified WT parasites. Primers used are as indicated in Figure 1, demonstrating a product across the 5’ and 3’ integration junction (indicated as 5’ int and 3’ int, respectively) as well as quantitative absence of the original locus (‘orig. loc.’). Absence of this band indicates that no WT parasites remained in the parasite population. KI, knock in cell line. For primer sequences see Supplementary file 1.

**Figure 5 – figure supplement 1.**
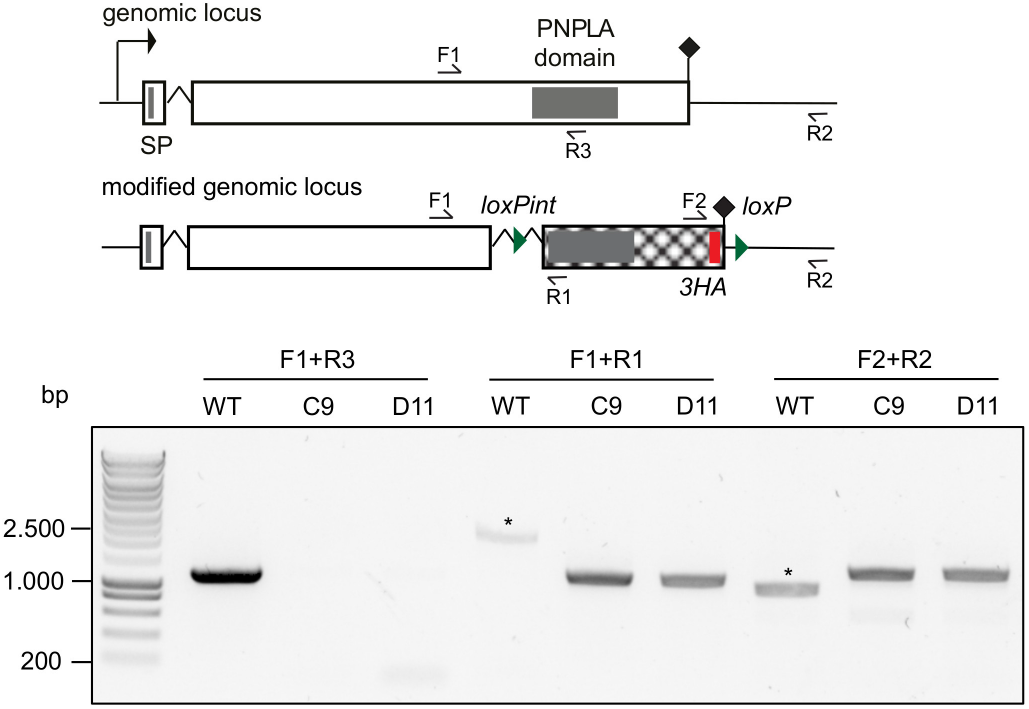
Integration PCR of PNPLA2:HA:loxPint parasites. Schematic of the *pnpla2* locus before and after CRISPR-Cas9-based gene editing is shown on top, while agarose gel electrophoresis of PCR products from unmodified WT and clonal modified parasite lines are displayed below. Primers used for confirming correct integration into the genome are indicated with arrows. Non-specific PCR products are marked with an asterisk. For primer sequences see Supplementary file 1.

**Supplementary file 1.** Oligonucleotides and other synthetic DNA used in this study.

**Supplementary file 2.** Lipidomic analysis of untreated and Rapa-treated PI-PLC-GFP-KS parasites.

**Supplementary file 3.** Details on the lipid standard used for lipidomic analysis.

